# Protocadherin-mediated cell repulsion controls the central topography and efferent projections of the abducens nucleus

**DOI:** 10.1101/283473

**Authors:** Kazuhide Asakawa, Koichi Kawakami

## Abstract

Cranial motor nuclei in the brainstem innervate diverse types of head and neck muscles. Failure in establishing these neuromuscular connections causes congenital cranial dysinnervation disorders (CCDDs) characterized by abnormal craniofacial movements. However, mechanisms that link cranial motor nuclei to target muscles are poorly understood at the molecular level. Here, we report that protocadherin-mediated repulsion mediates neuromuscular connection in the ocular motor system in zebrafish. We identify pools of abducens motor neurons that are topographically arranged according to soma size and convergently innervate a single muscle. Disruptions of Duane retraction syndrome-associated transcription factors reveal that these neurons require Mafba*/*MAFB, but not Sall4/SALL4, for differentiation. Furthermore, genetic perturbations of Pcdh17/Protocadherin-17 result in defective axon growth and soma clumping, thereby abolishing neuromuscular connectivity. Our results suggest that protocadherin-mediated repulsion forms the central topography and efferent projection pattern of the abducens nucleus following Mafba-dependent specification, and imply potential involvement of protocadherins in CCDD etiology.

## Introduction

Diverse movements of the head, face and eyes are regulated by cranial motor nuclei in the brainstem via complex efferent nerve pathways (Gilland and Baker, 2005; Guthrie, 2007). In adult vertebrates, cranial motor nuclei are typically organized in rostrocaudal columns, and include motor neurons originating from transverse hindbrain segments known as rhombomeres established during the embryonic period (Lumsden and Keynes, 1989; Orr, 1887; Vaage, 1969). For individual cranial motor neurons to connect with appropriate muscle fibers, not only specification but also subsequent axon growth and guidance must be regulated with remarkable precision. Concurrently, developing motor neurons with similar functions cluster together into a discrete nucleus, and neurons within nuclear subdivisions collectively projected onto peripheral target muscles often in a topographic manner. Such individual- and collective-level regulation of cranial motor neuron development has been intensively explored by analyzing causative genes for human congenital cranial dysinnervation disorders (CCDDs) and by investigating tractable cranial motor systems in experimental animal models (Astick et al., 2014; Chilton and Guthrie, 2016; Whitman and Engle, 2017). Nevertheless, these mechanisms remain incompletely understood.

Abducens motor neurons control abduction of the eyes by innervating the lateral rectus muscle. While the abducens nerve targets a single muscle, the abducens nucleus exhibits diverse distribution patterns in the hindbrain among vertebrate taxa. For instance, it is located in both rhombomere (r)5 and r6 in teleosts, reptiles and birds, but is restricted to r5 in mammals and to r6 in elasmobranchs. The abducens nucleus originating from a single rhombomere includes multiple types of motor neurons that are topographically distributed (Buttner-Ennever et al., 2001; Eberhorn et al., 2005; Eberhorn et al., 2006), and projects an efferent nerve consisting of structurally segregated sub-fibers (da Silva Costa et al., 2011; Peng et al., 2010). The lateral rectus muscle also contains structural subdivisions, the global (inner) and orbital (outer) layers with differential content of physiologically distinct myofibers, including those mediating tonic or phasic contraction (Morgan and Proske, 1984; Siebeck and Kruger, 1955; Spencer and Porter, 1988, 2006). Neither the individual-level nor collective-level connectivity between these subdivisions of abducens nucleus and lateral rectus muscle has been fully described, mainly due to the limited anatomical accessibility of the entire neuronal projection. The anatomy and physiological significance of the segmentally separated teleostean and avian abducens nucleus also remain poorly understood, despite intensive investigations (Cabrera et al., 1989; Gestrin and Sterling, 1977; Pastor et al., 1991; Sterling and Gestrin, 1975).

The specification of the abducens motor neurons relies on Hox-dependent positional values (Gaufo et al., 2003; Manzanares et al., 1997; Prince et al., 1998). Following specification, axons of abducens motor neurons are guided by extrinsic directional cues, including semaphorin and ephrin, through activation of downstream signaling pathways that modulate cytoskeletal dynamics (Ferrario et al., 2012) (Nugent et al., 2017). Consistent with these molecular mechanisms, the causative genetic loci for Duane retraction syndrome (DRS), a congenital type of strabismus, encode transcription factors for hindbrain patterning (*HOXA1* and *MAFB*) (Tischfield et al., 2005) (Park et al., 2016) and a Rac-GAP modulating cytoskeletal dynamics (*CHN1/*α2-chimaerin) (Miyake et al., 2008). In addition, DRS mutations are found in the gene encoding SALL4 transcription factor, but the contribution of *SALL4* to DRS pathology is unknown at the molecular level (Al-Baradie et al., 2002; Kohlhase et al., 2002). Currently, molecular mechanisms that spatially organize multiple types of abducens motor neurons within a nucleus and connect each motor neuron subtype to appropriate target myofibers are unknown. Cadherin superfamily cell adhesion molecules are differentially expressed in cranial motor nuclei (Astick et al., 2014) (Liu et al., 2015), and a specific type of cranial motor neuron can be mispositioned upon global misexpression of cadherin-20 in chick (Astick et al., 2014). However, whether and how cadherin superfamily genes contribute to formation of cranial neuromuscular connectivity is unknown.

The genetic amenability and translucency of zebrafish (*Danio rerio*) are uniquely suited for studying cellular and genetic bases of ocular motor pathway (Asakawa et al., 2012; Clark et al., 2013; Greaney et al., 2017; Higashijima et al., 2000; Kasprick et al., 2011; Lyons et al., 2010; Ma et al., 2014). Here, we identify pools of abducens motor neurons located in r5 and r6 using the *cis*-elements of the *mnr2b*/*mnx2b* homeobox gene. These abducens motor neurons are topographically arranged according to soma size along the dorsoventral axis, and require the DRS-associated hindbrain patterning transcription factor Mafba for differentiation. Furthermore, perturbations of Pcdh17/Protocadherin-17 by gene knockout and dominant-negative approaches result in the clumping of abducens motor neuron somas and axons, abrogating the central topography and axon growth. Based on these observations, we propose a novel mechanism for establishing collective-level cranial neuromuscular connectivity in which protocadherin-mediated repulsion promotes the formation of topographic organization, as well as its efferent projection, of the motor nucleus.

## Results

### Visualizing neuromuscular connections between the abducens nucleus and lateral rectus muscle in zebrafish

To visualize neuromuscular connections between the abducens nucleus and lateral rectus muscle, we first generated a transgenic zebrafish Gal4 driver for abducens motor neurons using a bacterial artificial chromosome (BAC) carrying the *hsp70l* promoter-linked Gal4FF (Asakawa et al., 2008) (hereafter, Gal4) in the *mnr2b*/*mnx2b* locus (Fig. 1A, B, Tg[mnr2b-hs:Gal4]). In Tg[mnr2b-hs:Gal4, UAS:enhanced green fluorescent protein (EGFP)] larvae, the EGFP-expressing cells were bilaterally aligned in both rhombomere(r)5 and r6 from 2 days post-fertilization (dpf) (Fig. 1C, D). In each of these rhombomere hemisegments, the EGFP-expressing cells were segregated dorsoventrally into two clusters (Fig. 1E, F). The nerve bundles emanating from these EGFP-expressing cell clusters extended ipsilaterally and terminated in the vicinity of the caudal eye globe, showing that the EGFP-expressing cells included abducens motor neurons.

**Figure 1.**
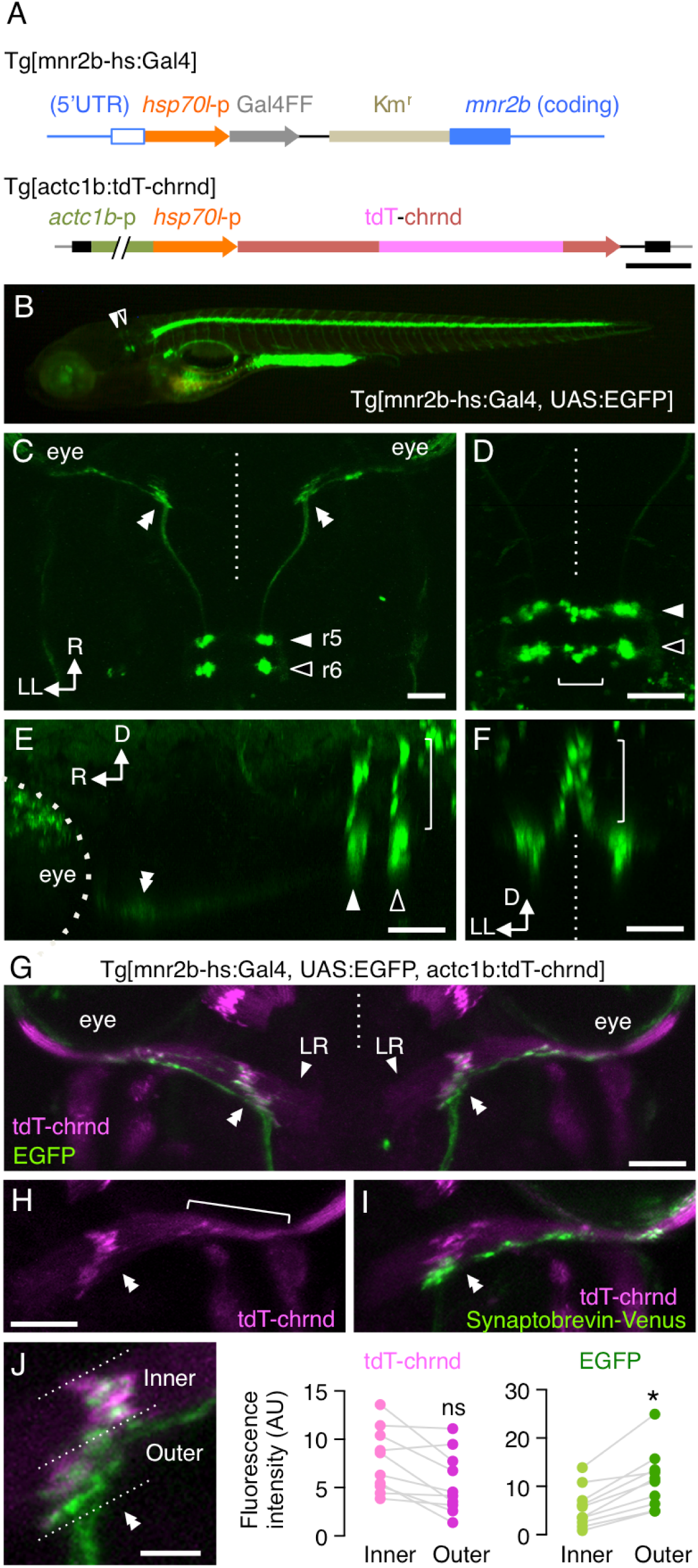
The neuromuscular connection between mnr2b-ABNs and the lateral rectus muscle. (A) Structure of Tg[mnr2b-hs:Gal4] and Tg[actc1b:tdT-chrnd]. (top) The *hsp70l* promoter-Gal4FF-polyA-Km^r^ cassette was inserted between the 5’ untranslated region (UTR) and the coding region of *mnr2b.* (bottom) tdT-chrnd is driven by the *hsp70l* promoter fused to the *actc1b* promoter. (B) Lateral view of Tg[mnr2b-hs:Gal4, UAS:EGFP] larvae at 5dpf. (C) Dorsal view of projection of the EGFP-labeled abducens motor axons. The dorsal EGFP-expressing cells, which are indicated with a bracket in D, are omitted for clarity of the projections. (D-F) The magnified views of EGFP cell clusters in C from the dorsal (D), lateral (E) and rear (F) directions. (G) Dorsal view of Tg[mnr2b-hs:Gal4, UAS:EGFP, actc1b:tdT-chrnd] larvae at 5 dpf. (H) Gathered (double arrowhead) and scattered (bracket) AChRs visualized by tdT-chrnd on the right lateral rectus muscle in G. (I) The right lateral rectus muscle of Tg[mnr2b-hs:Gal4, UAS:V2V, actc1b:tdT-chrnd] larvae at 5 dpf. (J) The fluorescence intensities of the gathered tdT-chrnd and associated EGFP signals were measured for each inner and outer halves of the muscle and subtracted by those of background signals in the adjacent muscle areas. N = 10 neuromuscular connections from 5 animals. *, p = 0.024. ns, not significant, p = 0.12. (unpaired *t* test). In G-J, rostral is to the top. The double arrowheads indicate the presumptive initial contact sites with the lateral rectus muscle. White and black arrowheads indicate the EGFP clusters in r5 and r6, respectively. Horizontal and vertical brackets indicate the cells in the dorsal cluster. Dashed lines indicate the midline. Bars indicate 500 base-pair (bp) in A, 50 µm in C, D, E, F, H, I, and 20 µm in J. R; rostral. D; dorsal. LL; left-lateral. LR; lateral rectus.

Next, to determine the efferent innervation pattern of these EGFP-expressing abducens motor neurons, we counterstained the lateral rectus muscle by contructing a transgene driving expression of td-Tomato-tagged acetylcholine receptor (AChR) δ subunit (tdT-chrnd) from the *actc1b* promoter (Fig. 1A, G). The tdT-chrnd signal revealed both gathered and scattered AChR clusters in the middle and lateral regions of the muscle, respectively, as well as the entire muscle with the weaker signal in the plasma membrane starting at 3 dpf onwards (Fig. 1H). The EGFP-labeled axon terminals were in close apposition to both the gathered and scattered AChRs (Fig. 1G) and distributed more densely in the outer muscle (i.e.. distal to the eye globe) than the inner muscle (proximal to the eye globe) (Fig. 1 J). The biased terminal distribution was coincided with the distribution of the presynaptic vesicle protein synaptobrevin/VAMP (Fig. 1I), suggesting that the EGFP-expressing abducens motor neurons primarily innervate the outer muscle. To examine the potential inner-outer muscle differentiation, we screened approximately one thousand Gal4 gene trap lines (Asakawa et al., 2013) and identified two showing distinct lateral rectus expression. Tg[gSAIzGFFM773B] line selectively labeled the outer muscle by trapping the troponin T type 1 gene (*tnnt1*) expressed in slow myofiber, while Tg[gSAIzGFFM1068A] line displayed broad Gal4 expression by trapping the troponin T type3 gene (*tnnt3b*) expressed in fast myofibers (SupFig. 1), suggesting that the outer muscle includes a characteristic of slow fibers, as is the case in primates (Khanna et al., 2004; Spencer and Porter, 2006). Taken together, we conclude that Tg[mnr2b-hs:Gal4] labels abducens motor neurons topographically organized along the dorsoventral and rostrocaudal axes, and that these neurons preferentially innervate the outer lateral rectus muscle including both slow and fast fibers. Henceforth, we refer to the abducens motor neurons labeled with Tg[mnr2b-hs:Gal4] as mnr2b-ABNs.

### mnr2b-ABNs form soma size topography independently of target myofiber type and birth order

The heterogeneous morphology of the axon terminals suggests that mnr2b-ABNs include multiple motor neuron subtypes (Fig. 2A). To investigate differences among individual mnr2-ABNs, we injected the UAS:EGFP plasmid into Tg[mnr2b-hs:Gal4] embryos at the one-cell-stage for sparse labeling of individual neurons. Out of approximately 900 injected larvae, we identified forty single somas of mnr2b-ABNs in r5 and r6, in addition to one descending interneuron (SupFig. 2). Of these, six located in the ventral hindbrain were the only labeled mnr2b-ABNs in a hemi-brain, and unambiguously revealed that the morphologies of axon terminals could be categorized into two main classes. The first class included mnr2b-ABNs with a terminal possessing linearly spaced boutons (“*en grappe*” terminals), likely innervating multiply innervated muscle fibers (MIFs) with slow fiber properties (Fig. 2B, C). The second class included mnr2b-ABNs with a single club-shaped terminal likely innervating singly innervated muscle fibers (or SIFs) with fast fiber properties (Fig. 2 D, E). These terminals accumulated in the putative central *en plaque* endplate zone (CPEZ) in the middle of the muscle as in higher vertebrates (Buttner-Ennever et al., 2001). The *en grappe-* and *en plaque* terminals were also visible when dorsal mnr2b-ABNs alone were labeled (Fig. 2F), although the terminals were thinner and somas were smaller compared to ventral mnr2b-ABNs (Fig. 2G). These observations suggest that the dorsoventral topography of mnr2b-ABNs reflects soma size rather than target myofiber type.

**Figure 2.**
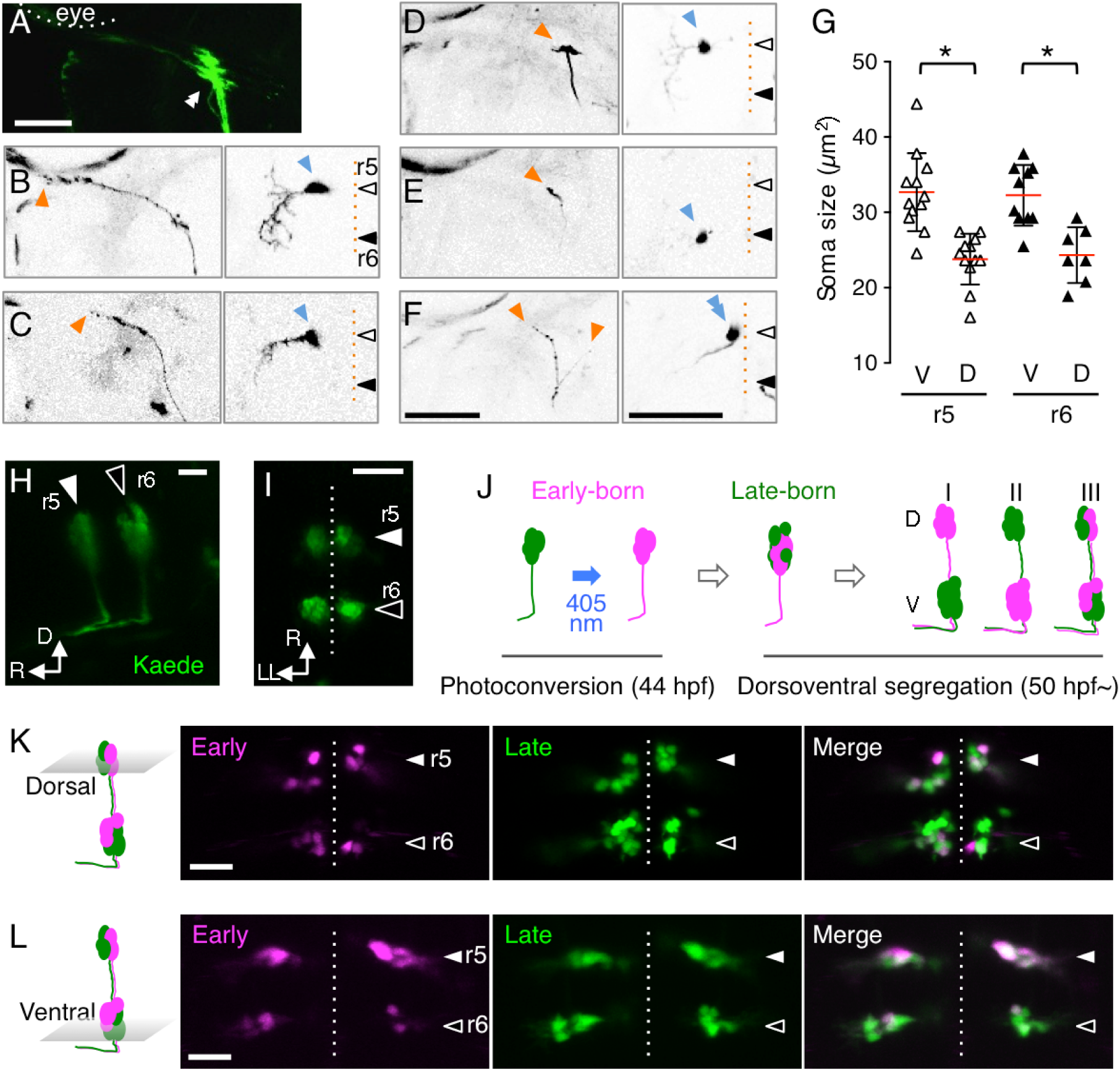
Soma size topography of mnr2b-ABNs. (A) Dorsal view of the terminal of the left abducens nerve of Tg[mnr2b-hs:Gal4, UAS:EGFP] larva at 5dpf. The double arrowheads indicate the presumptive initial contact sites with the lateral rectus muscle. (B-E) Axon terminals (left panels) and somas (right panels) of single abducens motor neurons. Arrowheads in orange and blue indicate tips of the axon terminal and somas, respectively. (F) Two neighboring abducens motor neurons are located on the ventral side of r5. The left and right terminals are *en grappe*- and *en plaque*- types, respectively. (G) The soma size of single cells in the dorsal and ventral clusters in r5 and r6. N = 40 cells from 22 animals. *, p < 0.05 (*t* test). (H, I) The lateral (H) and dorsal (I) views of mnr2b-ABNs in the Tg[mnr2b-hs:Gal4, UAS:Kaede] embryo at 44 hpf. (J) Birth order labeling of mnr2b-ABNs. Early-born mnr2b-ABNs are labeled in red by Kaede photoconversion at 44 hpf by irradiation of a laser with 405 nm wavelength. The segregation patterns of the early- (magenta) and late-born (green) populations are investigated at 72 hpf. Three possible segregation patterns are theoretically conceivable, where the early-born neurons are distributed in either of the dorsal (I) or ventral (II) cluster, or in both (III). D and V indicate dorsal and ventral, respectively. (K, L) Dorsal views of the dorsal (K) and ventral (L) area of r5 and r6 in Tg[mnr2b-hs:Gal4, UAS:Kaede] larvae at 72 hpf. The bars indicate 50 µm in A-F and 20 µm in H, I, K, L. White and black arrowheads indicate rhombomere identity. The vertical dashed lines indicate the midline. The rostral is to the top, unless otherwise indicated.

To determine whether this soma size topography is defined by the temporal order of neuronal differentiation, as suggested by *mnr2b*-labeled spinal motor neurons, which consist of early-born larger neurons located more dorsally and late-born smaller neurons located more ventrally (Asakawa et al., 2013). We discriminated early- from late-differentiating mnr2b-ABNs using the photoconvertible protein Kaede. In Tg[mnr2b-hs:Gal4, UAS:Kaede] larvae, mnr2b-ABNs first appeared as a single cell cluster in a hemisegment prior to dorsoventral segregation (Fig. 2H, I)(Asakawa et al., 2012). The early differentiating mnr2b-ABNs were marked with red fluorescence by photoconverted Kaede at 44 hours post-fertilization (hpf), and the subsequent distribution pattern along the dorsoventral axis was examined at 72 hpf (Fig. 2J). Unexpectedly, we found that the early differentiating mnr2b-ABNs were distributed in both the dorsal and ventral clusters (Fig. 2K, L), suggesting that larger and smaller mnr2b-ABNs concurrently differentiate before topographic segregation. Taken together, we conclude that mnr2b-ABNs in a rhombomere hemisegment are topographically arranged based on soma size rather than target fiber type and neuronal birth order.

### mnr2b-ABNs in r5 and r6 form convergent efferent projections

In the above experiments, we did not find any major differences in soma distribution pattern or developmental timing between mnr2b-ABNs in r5 and r6, but it remains uncertain whether mnr2b-ABNs in r5 and r6 show differences in efferent innervation patterns. To address this question, we developed a Gal4/Cre intersection approach to visualize mnr2b-ABNs in each of these rhombomeres by EGFP. We created two dual-color intersection UAS reporter lines. One reporter, Tg[UAS:loxRloxG], labeled r5 with EGFP via a ubiquitous Gal4 driver when the floxed mRFP1 was removed from r5 by the *egr2b*/Krox20-Cre transgene (Tg[egr2b-Cre])(Fig. 3A-C), while the other reporter, Tg[UAS:loxGloxR], labeled r6 with EGFP due to Cre-mediated removal of the floxed EGFP in r5 (Fig 3D, E).

**Figure 3.**
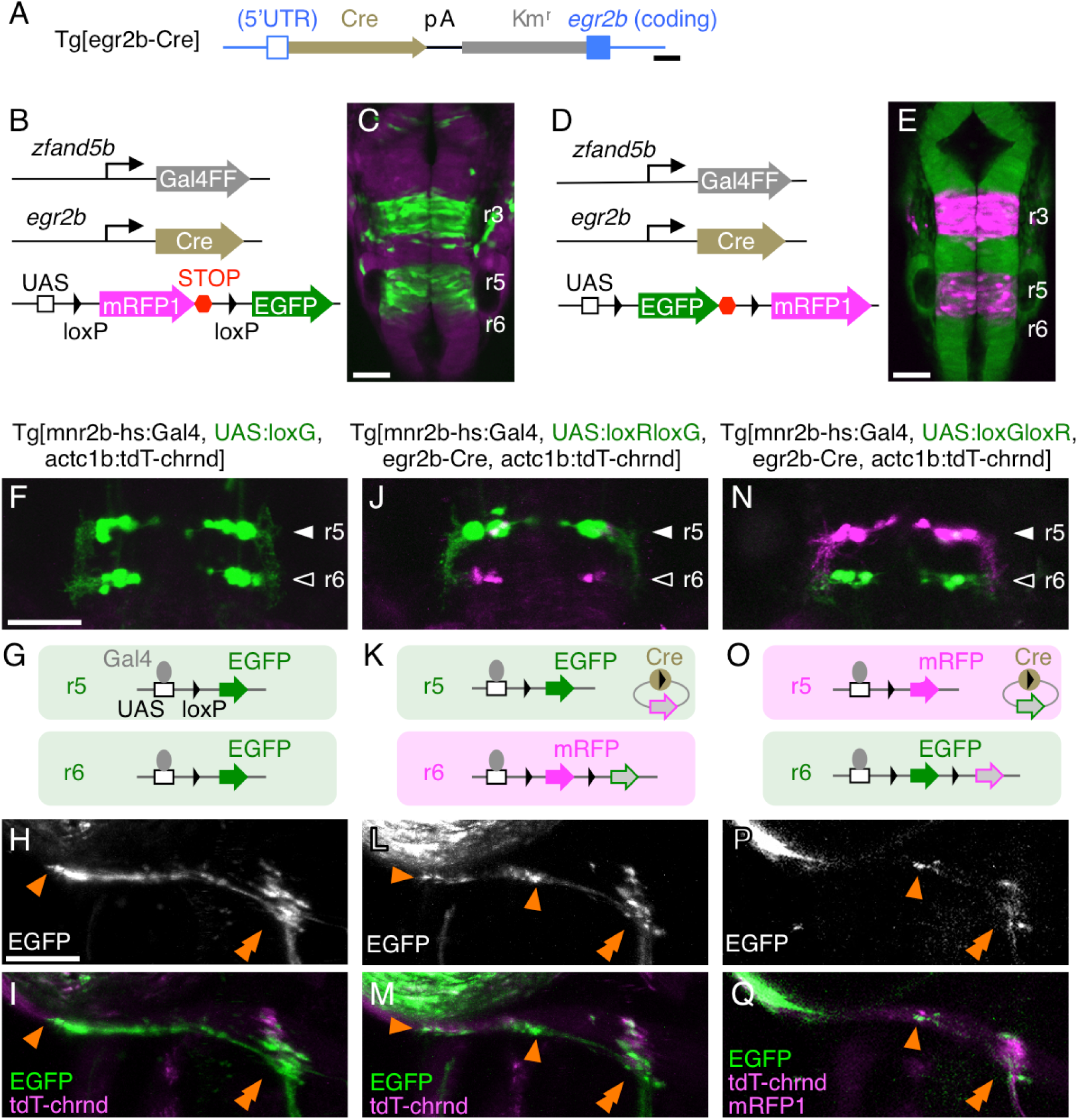
mnr2b-ABNs in r5 and r6 form convergent efferent projections. (A) Structure of Tg[egr2b-Cre]. The Cre-polyA-Km^r^ cassette was inserted between the 5’ untranslated region (UTR) and the coding region of *egr2b*. (B, C) EGFP expression in r5 in Tg[zfand5b-Gal4, egr2b-Cre, UAS:loxRloxG] embryo at 20 hpf. Tg[zfand5b-Gal4] is a ubiquitous Gal4 driver (Asakawa and Kawakami, 2009). (D, E) EGFP expression in r6 in Tg[zfand5b-Gal4, egr2b-Cre, UAS:loxGloxR] embryo. (F, J, N) Dorsal views of r5 and r6 in larvae at 5 dpf carrying the transgenes indicated above. (G, K, O) Schematic illustrations of Gal4- and Cre-mediated expression of EGFP and mRFP1 in r5 and r6. (H, I, L, M, P, Q) The axon terminals of mnr2b-ABNs innervating the left lateral rectus muscle. Tg[UAS:loxG] in F is created by excising the floxed mRFP1 from Tg[UAS:loxRloxG] in the germ line. The orange arrowheads indicate the terminal and distinct bouton structures of *en grappe* terminals. The double arrowheads show the CPEZ. In M, mRFP1 signal in mnr2b-ABN in r6 is below detection limit. The bars indicate 200 bp in A, and 50 µm in C, E, F and H.

We first labeled mnr2b-ABNs in r5 with EGFP by combining Tg[mnr2b-hs:Gal4], Tg[UAS:loxRloxG], Tg[egr2b-Cre], and Tg[actc1b:tdT-chrnd] transgenes (Fig. 3 J, K). In the quadruple transgenic fish, the axon terminals were detected both in the CPEZ and around the caudal end of the eye globes (Fig. 3L, M), indicating that the r5 subnucleus includes both *en grappe*- and *en plaque* type mnr2b-ABNs. This axon terminal distribution was quite similar to that of all mnr2b-ABNs in r5 and r6 (Fig. 3F-I). Next, we labeled mnr2b-ABNs in r6 with EGFP using the Tg[UAS:loxGloxR] reporter (Fig. 3N, O), and found that the r6 subnucleus also included mnr2b-ABNs with both *en grappe* and *en plaque* terminal morphologies (Fig. 3P, Q). Collectively, these results suggest that mnr2b-ABNs in r5 and r6 contain similar types of mnr2b-ABNs and form convergent efferent projections onto the lateral rectus muscle. The intersectional labeling also clearly revealed an elaborate dendritic network of mnr2b-ABNs in r5 emanating laterally and then trespassing into r6 (Fig. 3 J, N), implying mnr2b-ABNs in r5 and r6 have overlapping lateral dendritic fields.

### All mnr2b-ABNs require *mafba, but not sall4,* for specification

To elucidate the molecular mechanisms underlying the topographic organization and efferent projection patterns of mnr2b-ABNs, we investigated the effects of Duane retraction syndrome (DRS) gene knockout on mnr2b-ABN development. Haploinsufficient and dominant-negative mutations of the transcription factor gene *MAFB* cause DRS in humans (Park et al., 2016), and mutations of mouse Mafb (also known as Kreisler or Krml1) result in the absence or hypoplasia of abducens nerve, recapitulating DRS pathology (McKay et al., 1994; Park et al., 2016). To address how the zebrafish orthologue of *MAFB* (*mafba*/*valentino*) contributes to mnr2b-ABN development, we used a novel *mafba* mutant (*val*^M35A-Gal4^) in which its only exon was disrupted by the Gal4 gene trap insert Tg[gSAIzGFFM35A] (Fig. 4A) (Asakawa et al., 2013). As reported in the *val^-^* mutant (Ma et al., 2014; Moens et al., 1996), the homozygous *val*^M35A-Gal4^ embryos formed a vestigial r5 and r6 with diminished rostrocaudal length at 24 hpf (Fig. 4 B-E), suggesting that *val*^M35A-Gal4^ is a null allele. The enhancer trap line for *mnr2b*, Tg[mnr2b-GFP] (Asakawa et al., 2012), was crossed with *val*^M35A-Gal4^ to visualize mnr2b-ABNs expressing EGFP in a Gal4-independent manner (SupFig. 3). We found that not only the axons but also somas of mnr2b-ABNs were entirely absent in both r5 and r6 of *val*^M35A-Gal4^ homozygotes at 4-6 dpf (N = 5 animals, Fig. 4F, G). Furthermore, all mnr2b-ABNs expressed *olig2*, which is essential for abducens motor neurons specification, using Tg[olig2:egfp] marker (Zannino and Appel, 2009) (Fig. 5H-K). Collectively, these observations indicated that mnr2b-ABNs require *mafba* for specification and are descendants of *olig2*-expressing neural progenitors in both r5 and r6.

**Figure 4.**
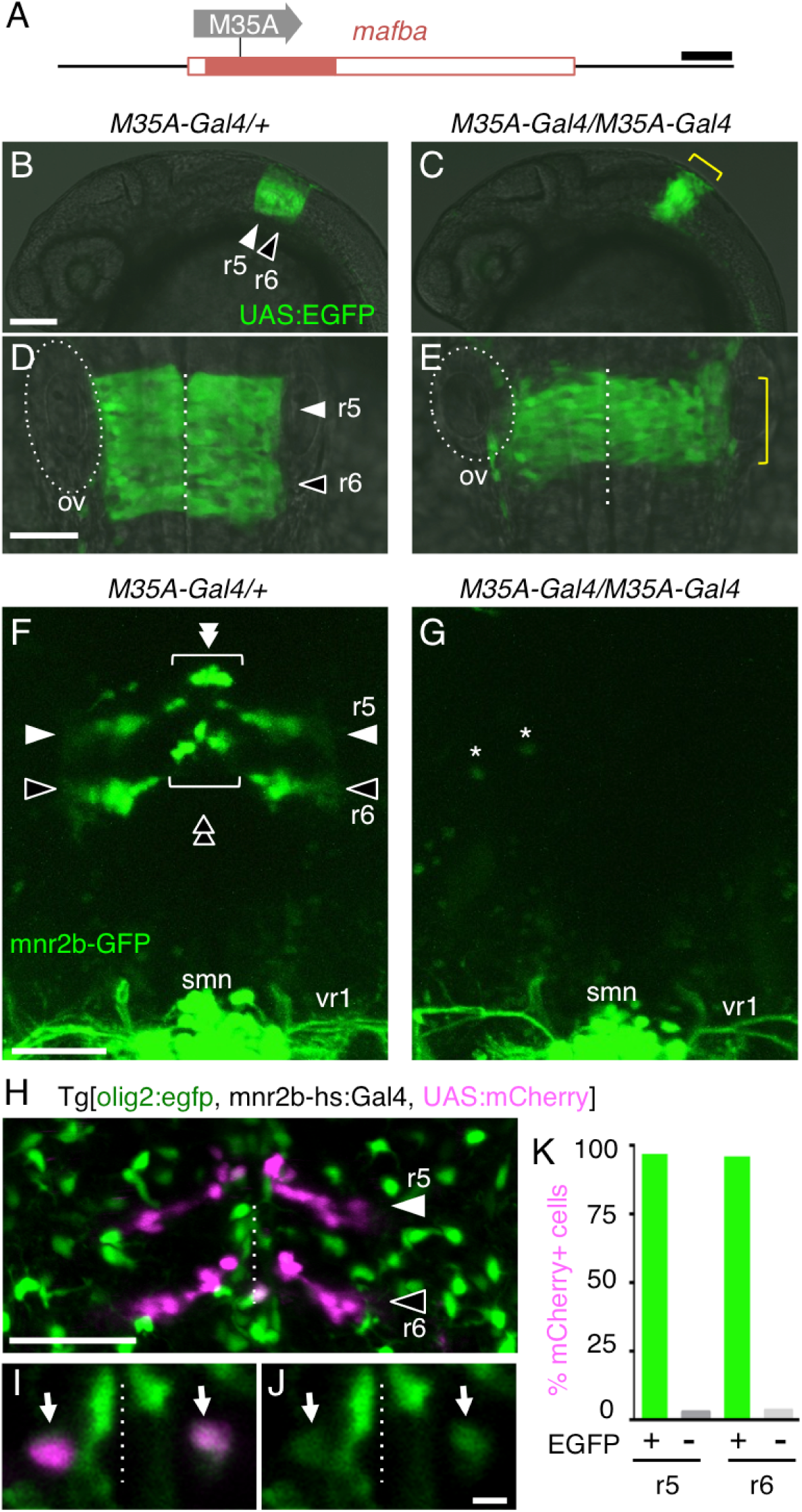
All mnr2b-ABNs require *mafba* for differentiation. (A) The *mafba*/*valentino* locus in Tg[gSAIzGFFM35A] line. Tg[gSAIzGFFM35A], indicated as M35A, is inserted in the coding exon of *mafba* (filled red box). (B-C) Lateral views of the rostral part (B, C) and dorsal views of r5 and r6 regions (D, E) of the Tg[*val^M35A^*/+, UAS:EGFP/+] and Tg[*val*^M35A-Gal4^/ *val*^M35A-Gal4^, UAS:EGFP/+] embryos at 24 hpf. Dashed lines and ovals show the midline and otic vesicle (ov), respectively. The yellow brackets in C and E indicate the vestigial r5 and r6. (F, G) Dorsal views of the hindbrain and first spinal segment of Tg[*val*^M35A-Gal4^/+, mnr2b-GFP/+] and Tg[*val*^M35A-Gal4^/ *val*^M35A-Gal4^, mnr2b-GFP/+] larvae at 80 hpf. In F, the arrowheads and brackets with double arrowheads indicate the dorsal and ventral mnr2b-ABNs, respectively. The white and black arrowheads/double arrowheads show mnr2b-ABNs in r5 and r6, respectively. Asterisks in G show EGFP-expressing cells unrelated to mnr2b-ABNs. smn; spinal motor neuron. vr1; ventral root from the first spinal segment. (H) Dorsal view of r5 and r6 in Tg[olig2:egfp, mnr2b-hs:Gal4, UAS:mCherry] at 3 dpf. (I, J) A single confocal section of the dorsal hindbrain in H. Arrows show mnr2b-ABNs expressing EGFP from Tg[olig2:egfp]. (K) Percentage of the mCherry-expressing cells that expressed EGFP. In total, 78 and 87 cells in r5 and r6, respectively, were examined in five larvae. The bars indicate 200 bp in A, 100µm in B, 50 µm in D-H, and 5 µm in I and J.

**Figure 5.**
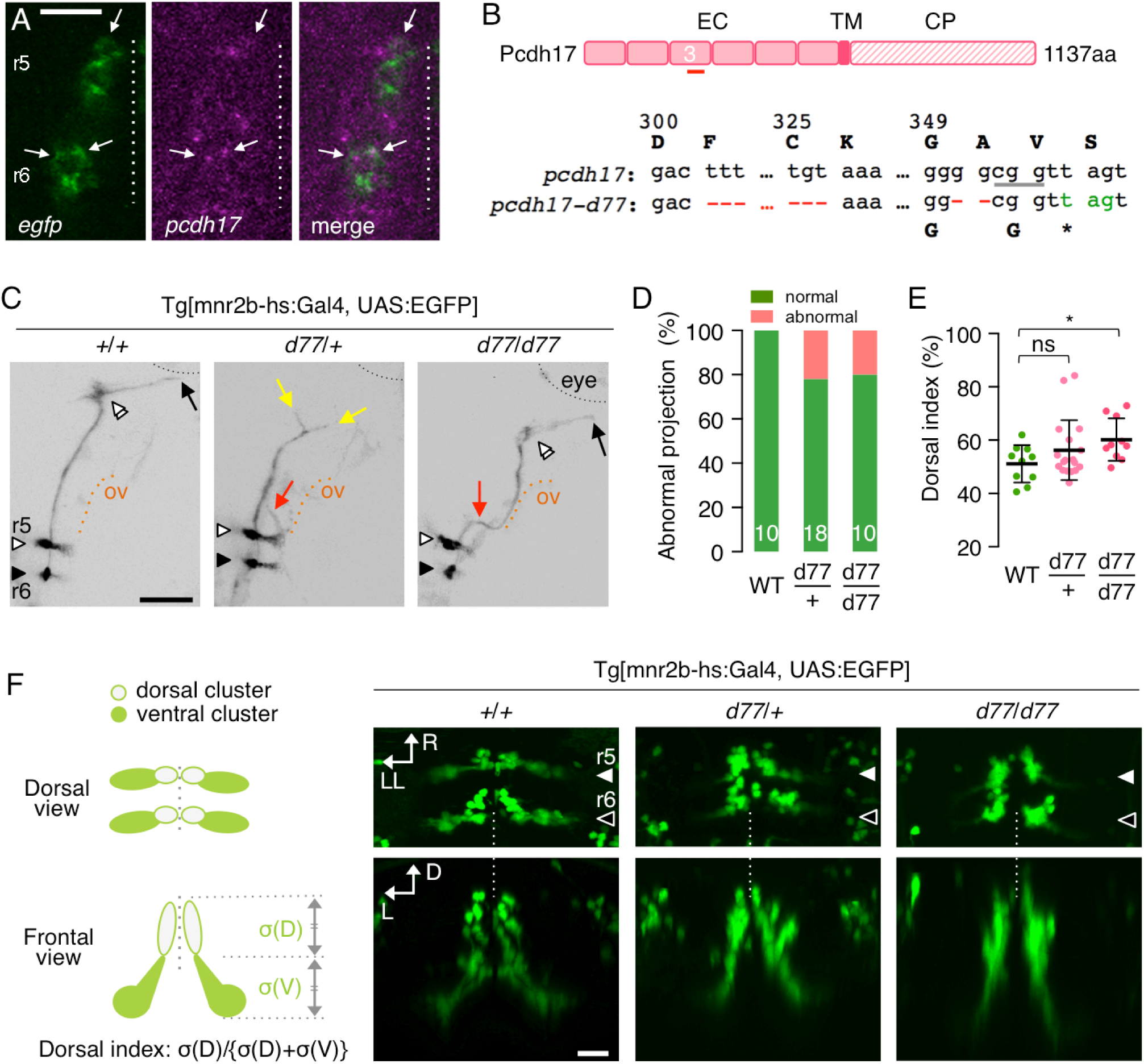
*pcdh17-d77* mutation causes defective axon growth and guidance, as well as soma topography formation. (A) Dorsal view of the left hemisegments of r5 and r6 of the Tg[mnr2b-hs:Gal4, UAS:EGFP] larva at 40 hpf. *EGFP* and *pcdh17* expression was detected by fluorescence *in situ* hybridization. Enhanced *pcdh17* expression is indicated by arrows. (B) The *pcdh17-d77* allele contains two deletions (red) in the extracellular domain 3 (EC3) coding sequence generating a premature stop codon (green). The upper cases represent amino acids encoded by the codons, and the numbers above indicate amino acid positions. The underlined cgg indicates the protospacer adjacent motif (PAM). EC, TM, and CP stand for extracellular domain, transmembrane domain and cytoplasmic domain, respectively. (C) The axon growth (yellow arrows) and pathfinding (red arrows) defects of the abducens nerve in Tg[mnr2b-hs:Gal4, UAS:EGFP] larvae carrying the heterozygous (middle) and homozygous (right) *pcdh17-d77* mutation at 3 dpf. The black and orange dashed lines demarcate the eye and otic vesicle (ov), respectively. (D) Frequency of abnormal axonal projection shown in E observed in wild type (WT), heterozygous (d77/+) and homozygous (d77/d77) pcdh17-d77 mutant larvae. (E) Distribution of mnr2b-ABN somas is dorsally shifted in *pcdh17-d77* mutants. The dorsal index is defined by the ratio of the area of mnr2b-ABN somas in the dorsal half of the nucleus (σ(D)) to that of all somas (σ(D)+σ(V)) in an rostrocaudally projected image as shown in F. ns, not statistically significant (p = 0.15, unpaired t test). *, p = 0.0144. (F) Schematic illustrations of normal nuclear organization mnr2b-ABNs (left) and dorsal and frontal views of mnr2b-ABN somas at 3 dpf. The dashed lines indicate the midline. The bars indicate 20 µm in A and F, and 50 µm in C. The white and black arrowheads show mnr2b-ABNs in r5 and r6, respectively. R; rostral. D; dorsal. L; lateral. LL; left-lateral.

Haploinsufficiency of SALL4 transcription factor causes a specific type of DRS characterized by abducens nerve hypoplasia and radial ray defects (also known as Okihiro syndrome/Duane-radial ray syndrome) (Al-Baradie et al., 2002; Demer et al., 2007; Kohlhase et al., 2002). To investigate functions of the zebrafish orthologue of *SALL4* (*sall4*) in mnr2b-ABN development, we generated a frame shift mutation in the middle region of *sall4* (*sall4-d11*), the region where causative nucleotide insertions and deletions have been found in Okihiro syndrome patients, using by using CRISPR-Cas9 method (SupFig. 4A, B). The homozygous *sall4-d11* larvae were lethal, and exhibited morphological abnormalities, including cardiac and eye edema, smaller eye, shortened bodies (SupFig. 4C, D). To our surprise, however, *sall4-d11* homozygotes developed normal mnr2b-ABN clusters in r5 and r6 (SupFig. 4 F-J), as well as neuromuscular connection between mnr2b-ABNs and the lateral rectus muscle at 4-6 dpf (SupFig. 4E, H), indicating that *sall4* is not involved in the development of mnr2b-ABNs in zebrafish.

### *pcdh17* knockout causes abnormal axonal extension and disrupts the nuclear topography of mnr2b-ABNs

The cell-adhesion molecule Pcdh17/Protochadherin-17 (encoded by the *pcdh17* gene) is expressed in the abducens nucleus of adult zebrafish (Liu et al., 2015), and its mouse orthologue Pcdh17 regulates the axonal extension of amygdala neurons (Hayashi et al., 2014) and synaptogenesis in corticobasal ganglia circuits (Hoshina et al., 2013). In zebrafish, *pcdh17* is expressed as early as at 1.5−6 hpf and expression becomes enhanced in specific tissues, including the hindbrain, during larval stages (Biswas and Jontes, 2009; Liu et al., 2009). Therefore, we hypothesized that Pcdh17 may play a key role in axon projection of mnr2b-ABNs. By double-fluorescent *in situ* hybridization, we detected enhanced *pcdh17* expression in a subset of mnr2b-ABNs in r5 and r6 prior to their dorsoventral segregation as well as weak widespread expression across the hindbrain at 46 hpf (Fig. 5A). Using the CRISPR-Cas9 method, we then created a *pcdh17* mutant encoding a truncated Pcdh17 protein due to two neighboring deletions (75 and 2 base-pairs) in the middle of the extracellular domain (*pcdh17-d77*, Fig. 5B). We found that the *pcdh17-d77* mutation exhibited maternal effects because, among the offspring of *pcdh17-d77* incrosses, 31 % of embryos including *pcdh17-d77* homozygotes, heterozygotes and wild type siblings, displayed early developmental abnormalities, such as gastrulation defect (SupFig. 5). Moreover, 65% of the embryos that did not show such early defects failed to inflate swim bladder on time on 5 dpf with an occasional formation of cardiac edema. Similar developmental abnormalities were observed among the offspring from *pcdh17-d77* female outcrosses and, to a lesser extent, from *pcdh17-d77* male outcrosses, suggesting that both maternal and zygotic Pcdh17 are required for normal development. In *pcdh17-d77* homozygous as well as heterozygous larvae, we observed reduced axon growth and misguided axon projection of mnr2b-ABNs in approximately 20% of nerves at 3 dpf (Fig. 5C, D). Furthermore, the distribution pattern of mnr2b-ABN somas was shifted significantly to the dorsal side in *pcdh17-d77* homozygous larvae at 3 dpf (Fig. 5E, F). These observations indicate that *pcdh17* is involved in the axon growth and nuclear formation of mnr2b-ABNs, although the earlier and global effect of the *pcdh17-d77* mutation precluded a rigorous evaluation of *pcdh17* expressed within mnr2b-ABNs.

### Protocadherin-mediated repulsion regulates the topographic organization and axon projection of mnr2b-ABNs

In order to investigate cell-autonomous functions of Pcdh17 in mnr2b-ABNs, we then used the Gal4/UAS system to express a dominant-negative Pcdh17 mutant lacking the cytoplasmic domain, which normally serves to mediate signaling with cytoskeletal regulators (Pcdh17-ΔCP) (Hayashi et al., 2014) (Fig. 6A). A plasmid carrying mRFP1-tagged *pcdh17-ΔCP* (*pcdh17-ΔCP-mRFP1*) and EGFP on either side of UAS (Fig. 6A) was injected into Tg[mnr2b-hs:Gal4] embryos to label mnr2b-ABNs expressing *pcdh17-ΔCP-mRFP1* with EGFP. We found that mnr2b-ABN axons expressing Pcdh17-ΔCP-mRFP1 formed clumps either at the shaft or terminal region in approximately 40% of nerves (Fig. 6C, D, 7/18 nerves, 9 animals), while this clumping phenotype was not observed in axons expressing the full-length Pcdh17 (*pcdh17-FL-mRFP1*) (Fig. 6B, 0/9 nerves, 6 animals), suggesting that Pcdh17 is cell-autonomously required for axon growth of mnr2b-ABNs.

**Figure 6.**
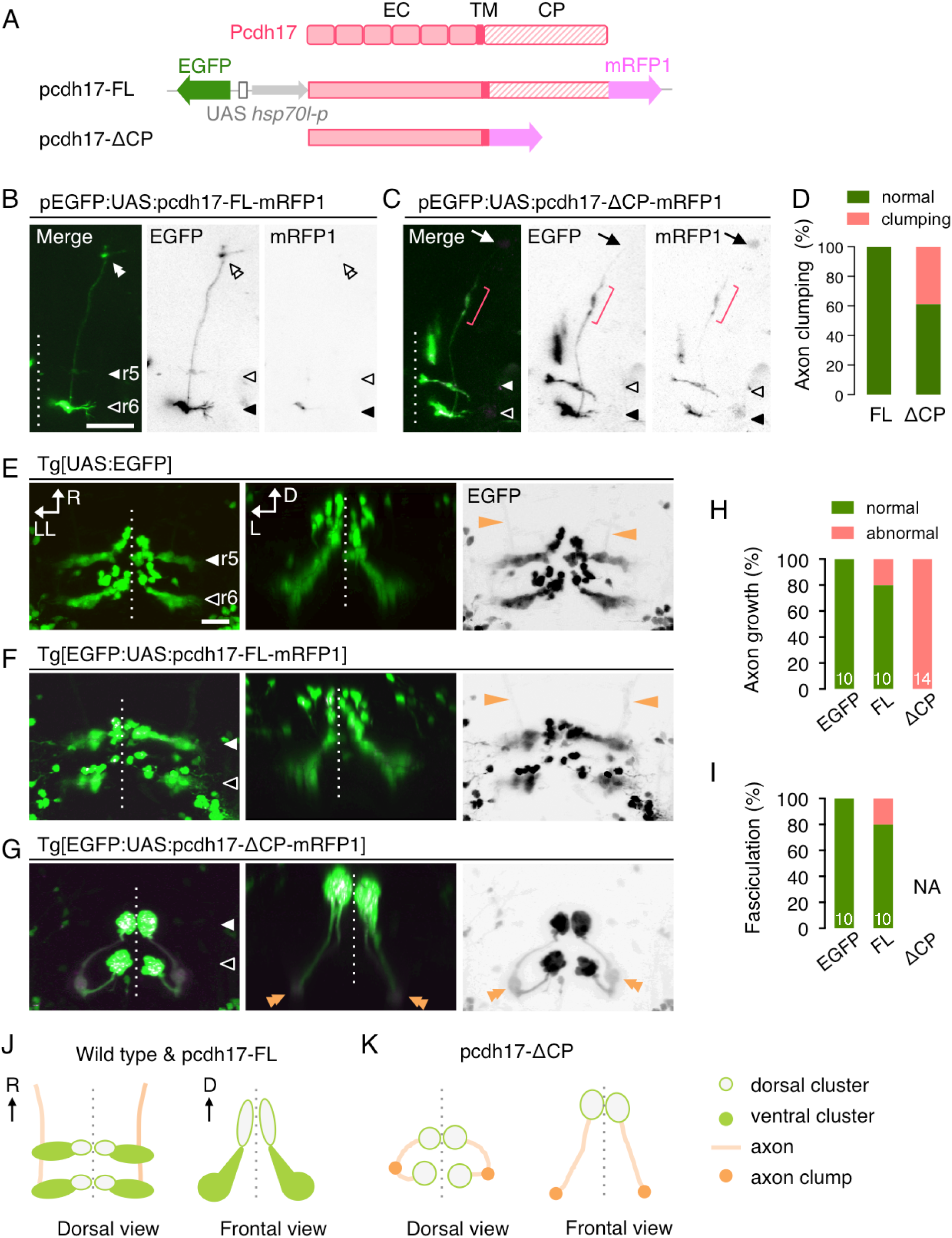
Dominant negative Pcdh17 disrupts nuclear topography and axon growth of mnr2b-ABNs. (A) (top) Schematic drawing of Pcdh17 protein. EC, TM, and CP stand for extracellular domain, transmembrane domain and cytoplasmic domain, respectively. (bottom) UAS constructs for pcdh17-FL and pcdh17-ΔCP induction. EGFP and the C-terminally mRFP1-tagged *pcdh17* genes were place on each side of 4xUAS. The *hsp70l* promoter was used as a basal promoter for the pcdh17 genes. (B, C) Dorsal views of the right brain hemispheres of Tg[mnr2b-hs:Gal4] larvae injected with the plasmid indicated as above. The double arrowheads indicate the presumptive initial contact sites with the lateral rectus muscle. Nerve terminal is indicated by arrow. Axon clump is indicated with a red bracket. (D) Frequency of the axon clumping occurring in the mnr2b-ABNs labeled with the plasmids described in B. For pcdh17-FL and pcdh17-ΔCP, 9 nerves from 6 animals and 18 nerves from 11 animals were examined, respectively (E, F, G) Dorsal (left, right) and rear (middle) views of r5 and r6 in 3 dpf larvae carrying Tg[mnr2b-hs:Gal4] and the transgene indicated as above. The orange arrows and double arrowheads indicate the normal and clumped axons. (H) Frequency of abnormal axon growth. (I) Frequency of abnormal fasciculation. NA, not applicable due to severe axon growth defect. (J, K) Schematic illustrations of normal (J, Wild type and pcdh17-FL) and abnormal (K, pcdh17-ΔCP) nuclear formation and axon growth of mnr2b-ABNs. The dashed lines indicate the midline. The bars indicate 50 µm in B and C, and 20 µm in E-G. The white and black arrowheads show mnr2b-ABNs in r5 and r6, respectively. R; rostral. D; dorsal. L; lateral. LL; left-lateral.

Axonal clumping is indicative of loss of self-avoidance mediated by protocadherin homophilic interactions (Hoshina et al., 2013; Mountoufaris et al., 2017). We reasoned that if protocadherin-mediated self-avoidance were required for growth of mnr2b-ABN axons, then the whole axon bundle would clump due to persistent homophilic interactions when *pcdh17-ΔCP-mRFP1* was expressed in all mnr2b-ABNs. To test this possibility, we generated a transgenic zebrafish line carrying a stable transgene from which *pcdh17-ΔCP-mRFP1* and EGFP were simultaneously inducible by Gal4 (Fig. 6A). Strikingly, when *pcdh17-ΔCP* was expressed, the EGFP-labeled axons emanating from r5 and r6 were heavily clumped together around the ventral side of r5/r6 boundary and rostral extension was completely inhibited (Fig. 6G, H). Moreover, the somas of mnr2b-ABNs expressing *pcdh17-ΔCP-mRFP1* were tightly clustered on the dorsal side of the hindbrain and rarely observed on the ventral side. This clumping phenotype was not observed when *pcdh17-FL-mRFP1* was overexpressed from a transgene (Fig. 6F), although misguided axon projections and defasciculation were observed at a low frequency (Fig. 6I, SupFig. 6). The soma distribution of mnr2b-ABNs remained unaffected by overexpression of *pcdh17-FL-mRFP1* (Fig. 6E, F, G). The clumping of somas and axons caused by *pcdh17-ΔCP* expression required membrane targeting because deletion of the transmembrane domain from *pcdh17-ΔCP* (*pcdh17-ΔCPT*) completely abolished the clumping effect (SupFig. 7). Taken together, these observations suggest that protocadherin-mediated repulsion is required not only for axon growth but also for migration of abducens motor neurons. Intriguingly, induction of *pcdh17-ΔCP-mRFP1* expression in *mnr2b*-labeled spinal motor neurons resulted in similar clumping of axons and somas, and disrupted the normal dorsal localization of larger neurons in the spinal cord (SupFig. 8). This implies that protocadherin-mediated repulsion commonly operates in somatic motor neurons to promote axon growth and formation of soma topography.

## Discussion

Our analyses revealed that the abducens nerve pathway of zebrafish consists of cellular machineries conserved in higher vertebrates (Fig. 7A). These include motor neurons with *en grappe* and *en plaque* axon terminals, which most likely reflect their target myofiber types, MIFs and SIFs, respectively. The lateral rectus muscle also contains sub-structures identifiable in higher vertebrates, including the longitudinal inner/outer compartments characterize by distinct proportions of fast and slow myofibers and typical endplates accumulation, such as CPEZ (Spencer and Porter, 2006). However, it was unclear how these motor neurons are spatially arranged and connected to sub-compartments of the target muscle given that the abducens nucleus is segmentally iterated in teleosts. Single-cell, birth order, and Gal4/Cre intersectional labeling approaches indicated that mnr2b-ABNs in r5 and r6 are similar in neuronal type, differentiation timing, and efferent innervation pattern. In addition, the lateral dendritic fields of r5 and r6 mnr2b-ABNs are shared beyond their rhombomeric borders, suggesting that these abducens nuclei may receive common premotor inputs. Therefore, we conclude that the segmentally iterated teleostean abducens nuclei contain anatomically and functionally homologous motor neurons (Pastor et al., 1991) (Cabrera et al., 1992). A potential advantage of having such a homologous set of neurons is that the dynamic range of the lateral rectus muscle, and hence that of lateral eye movement, could be expanded by an increase in recruitable motor units (Fuchs and Luschei, 1970), which may be beneficial for survival of lateral-eyed animals such as fish. Our results, however, do not exclude the possibility that the projections from abducens nuclei in r5 and r6 are topographic within the inner/outer muscle compartments, since the positional difference between these abducens subnuclei may cause a temporal difference in the onset of innervation, potentially generating a local innervation topography.

**Figure 7.**
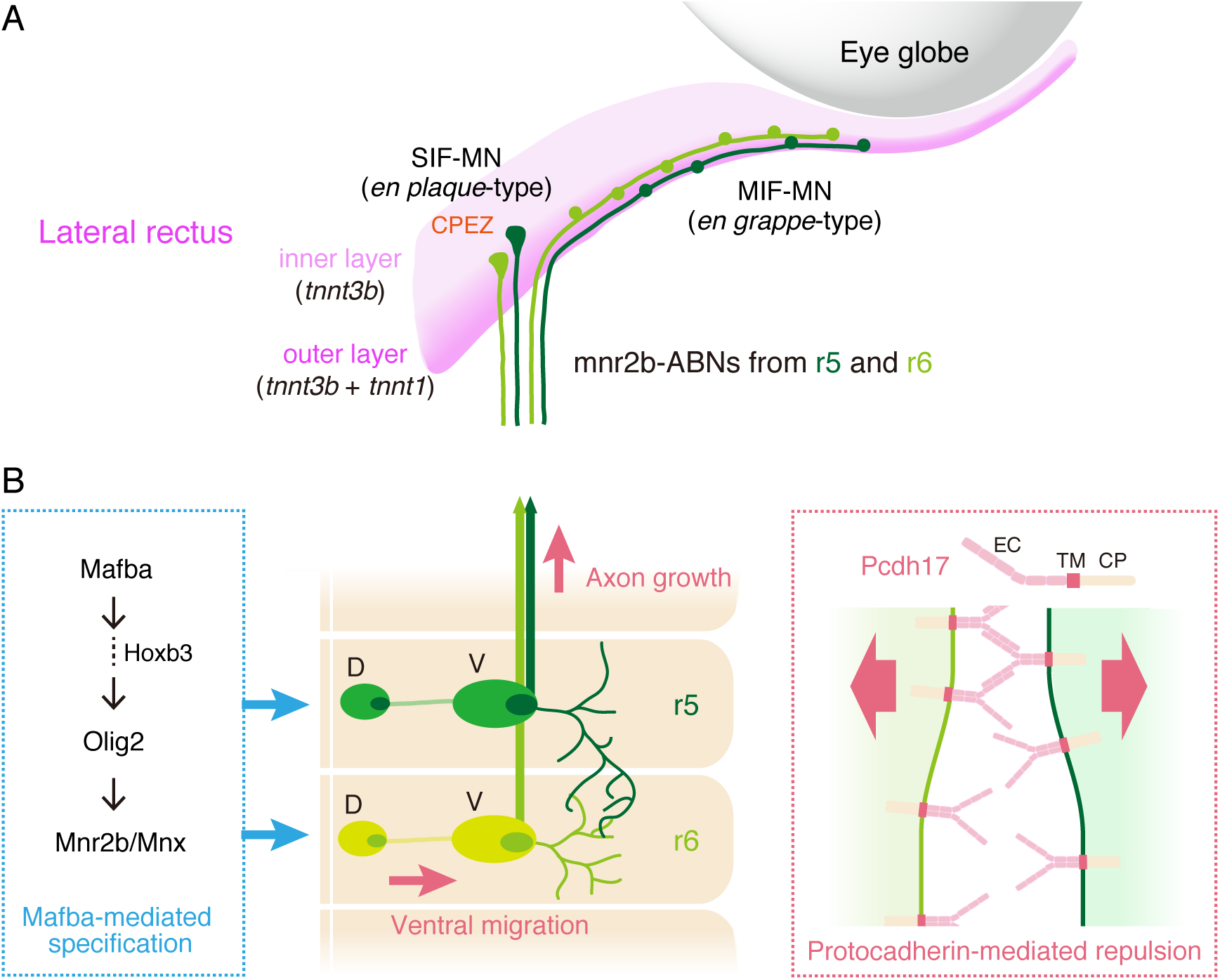
Architecture and developmental mechanism of the neuromuscular connection between mnr2b-ABNs and the lateral rectus muscle. (A) Schematic diagram of the lateral rectus muscle attached to the right eye globe. mnr2b-ABNs with *en plaque*-type terminal (SIF-innervating motor neurons, SIF-MN) innervates the CPEZ in the lateral rectus muscle, while ones with *en grappe*-type terminals (MIF-innervating motor neuron, MIF-MN) the outer lateral rectus muscle. The outer layer of the muscle contains SIF expressing the fast muscle-specific troponin T type3 (*tnnt3b*) and MIFs expressing the slow muscle-specific troponin T type 1 (*tnnt1*). mnr2b-ABNs originating from r5 and r6 form convergent projections to the lateral rectus muscle. (B) Schematic drawing of mnr2b-ABNs in the right hemisegment of r5 and r6. In each of r5 and r6, mnr2b-ABNs with smaller and larger somas are clustered dorsally and ventrally, respectively. The lateral dendrites of mnr2b-ABNs in r5 and r6 overlap each other. In both r5 and r6, mnr2b-ABNs require the Olig2-Mnr2b/Mnx cascade for differentiation (left). The Olig2-Mnr2b/Mnx pathway is activated by Mafba, possibly via Hoxb3. Protochadherin-mediated repulsive force (right) supports ventral migration and axon growth of mnr2b-ABNs.

In zebrafish, the genetic mechanisms underlying abducens motor neuron development are contentious as two conflicting conclusions were drawn from the *val^-^*/*mafba* mutant line. Our conclusion that Mafba is essential for the differentiation of mnr2b-ABNs (Fig. 7B) is in accordance with the observation that the abducens nucleus is rare or completely absent in the *val^-^* mutant (Moens et al., 1996), but inconsistent with the finding that the abducens nucleus exists in the *val^-^* mutant and is likely to have similar size as the wild type (Ma et al., 2014). We presume that the discrepancy may stem from the *olig2* enhancer trap line employed as an abducens nerve marker (Ma et al., 2014). It remains to be determined whether the labeled nerve bundle in the *val^-^* background originates solely from the abducens nucleus, given that the mutation in Mafba or Mafba-regulated genes can lead to ectopic neuronal differentiation (Gaufo et al., 2003) and ectopic muscle innervation (Park et al., 2016). Considering that virtually all mnr2b-ABNs were the descendants of progenitors expressing *olig2*+, which is essential for abducens motor neuron differentiation (Zannino and Appel, 2009), we propose that Mafba-Olig2-Mnx pathway serves as a common genetic cascade necessary for abducens motor neuron differentiation across vertebrates (Fig. 6B)(Lu et al., 2002; McKay et al., 1997; Thaler et al., 1999; Zhou and Anderson, 2002). Robust activation of *hoxb3/Hoxb3* appears necessary for development of the abducens nucleus as *hoxb3/Hoxb3* upregulation occurs in r5 and r6 of zebrafish but only in r5 of mice (Gaufo et al., 2003; Manzanares et al., 1997; Prince et al., 1998). Considering that Mafba/MAFB, which can directly activate *hoxb3/Hoxb3* (Manzanares et al., 1997), is expressed in r5 and r6 of both zebrafish and mice, the lack of an abducens nucleus in r6 of higher vertebrates may have been caused by extinction of Mafba-mediated *Hoxb3* activation specifically in r6 by an as yet unknown mechanism (Manzanares et al., 1997).

The abducens nucleus of zebrafish is topographically organized along the dorsoventral axis. This topography is based on soma size, not on target myofiber type or birth order, highlighting similarities and differences between abducens and spinal motor neurons, both of which are categorized as somatic motor neurons. From a functional point of view, soma size topography of the spinal motor column, where large and early-born neurons are located more dorsally, matches recruitment order during locomotion (McLean et al., 2007; Menelaou et al., 2014) (Ampatzis et al., 2014). How the soma size topography of abducens motor neurons relates to recruitment pattern for control of eye position and velocity is an important question to be addressed in the future (Robinson, 1970). From a developmental perspective, the soma clumping of the abducens and spinal motor neurons by Pcdh17-ΔCP expression argues that Pcdh17-dependent migration of larger motor neurons is a major driving force for the topographic soma distribution. These clumping phenotypes are likely due to loss of self-avoidance mediated by the homophilic interactions between Pcdh17 molecules (Hoshina et al., 2013), as observed when protocadherin function is compromised (Hayashi et al., 2014) (Lefebvre et al., 2012) (Mountoufaris et al., 2017). Thus, considering the everted configuration of the teleost brain, we propose that protocadherin-mediated repulsion promotes migration of larger motor neurons away from the ventricle and spinal canal (i.e., from the apical side of the hindbrain and spinal cord, respectively)(Holmgren, 1922). In addition to soma clumping, axon clumping of abducens and spinal motor neurons expressing Pcdh17-ΔCP suggests that axon growth is also regulated by protocadherin-dependent repulsion (Fig. 7B). Such repulsive forces among motor neurons may help maintain appropriate positional relationships in both the nucleus and nerve bundle, while still allowing individual neurons to respond to extrinsic directional axon guidance cues. Although we focused on *pcdh17* in this study, multiple cadherin superfamily molecules present on the cell surface could also be involved in nuclear topography formation and axon growth of abducens motor neurons. In fact, protocadherin *pcdh19* is also expressed in the zebrafish abducens nucleus (Liu et al., 2015). Functional redundancy of protocadherins may explain why the *pcdh17* knockout produced milder phenotypes in axon growth and soma topography than Pcdh17-ΔCP overexpression.

In principle, any gene mutation compromising abducens motor neuron development may contribute to eye movement disorders in humans. Indeed, mutations in *MAFB* (Park et al., 2016) and *HOXA1* (Tischfield et al., 2005), genes required for specification of abducens motor neurons, have been identified in DRS patients. Alternatively, the DRS-associated transcription factor *SALL4*/*sall4* is dispensable for abducens motor neuron specification in mice (Sakaki-Yumoto et al., 2006) and zebrafish, implying a human-specific function of *SALL4* in abducens motor neuron development. Our finding that the perturbation of *pcdh17* disrupted soma topography and axon growth of abducens motor neurons predicts that protocadherins are potential genetic candidates for human eye movement disorders. It is worth noting that the mouse *Pcdh17* has been identified as a major protocadherin in the MafB-expressing spiral ganglion cells, but not in closely related vestibular ganglion cells lacking MafB expression (Lu et al., 2011), raising the possibility that Pcdh17-mediated cell repulsion is an important cellular outcome of MafB-mediated transcriptional program in health and DRS pathology. Although pathological link between protocadherins and CCDDs remains to be proven, mutations in protocadherin genes could possibly contribute to CCDDs, given the distinct expression profiles of cadherin superfamily cell adhesion molecules in various cranial motor nuclei (Astick et al., 2014). Last, the ocular neuromuscular pathways in humans display selective vulnerability or resistance to certain types of muscular dystrophies and motor neuron diseases (Kaminski et al., 2002; Yu Wai Man et al., 2005), but mechanisms are relatively unexplored due to restricted tissue accessibility. We envision that the optically and genetically accessible abducens neuromuscular connection examined here could be of great help for interrogating pathologies of such diseases, as well.

## Acknowledgements

Authors would like to thank Kawakami lab members for valuable discussions, Dr Appel B for providing Tg[olig2:egfp] line, Miwako Okamura for helping UAS:loxRloxG construction, and Ito, M. Suzuki, M. Mizushina, N. Mouri and Y. Kanebako for fish maintenance. This work was supported by JSPS KAKENHI Grant numbers 23241063 (KK), 22700349 (KA), MEXT KAKENHI Grant number 23115720 (KA), The Uehara Memorial Foundation (KA), The Kao Foundation for Arts and Sciences (KA), The Mitsubishi Foundation (KA), Daiichi-Sankyo Foundation of Life Science (KA) and the National BioResource Project from the Ministry of education, Culture, Sports, Science and Technology of Japan (KK).

## Author contributions

KA conceived the research, designed and performed the experiments, analyzed the data. KA and KK performed gene trap screening. KA and KK wrote the manuscript.

## Competing interests

The authors declare no conflict of interest associated with this manuscript.

## STAR Methods

### Fish

This study was carried out in accordance with the Guide for the Care and Use of Laboratory Animals of the Institutional Animal Care and Use Committee (IACUC, approval identification number 24-2) of the National Institute of Genetics (NIG, Japan), which has an Animal Welfare Assurance on file (assurance number A5561-01) at the Office of Laboratory Animal Welfare of the National Institutes of Health (NIH, USA). The zebrafish larvae analyzed in this study were obtained by crossing the parental fish of either sex carrying appropriate transgenes, and the sex difference did not affect the results.

### Transgenic lines

Tg[mnr2b-hs:Gal4] was constructed as described in (Asakawa et al., 2013) except that the *hsp70l* promoter (650 bp)-Ga4FF-polyA-Km^r^ cassette was introduced downstream of the *mnr2b* 5’UTR in the *mnr2b*-BAC DNA (CH211-172N16, BACPAC Resources Center). Tg[mnr2b-hs:Gal4] line was chosen from five independent Tg[mnr2b-hs:Gal4] inserts, all of which gave rise to a similar Gal4 expression pattern as judged by UAS:EGFP reporter expression. Tg[actc1b:tdT-chrnd] was constructed by fusing the *chrnd* (GenBank accession number, AB120372) internally tagged with td-Tomato, as described in on the YFP-tagging of Chrnd (Epley et al., 2008), to the *hsp70l* and *actc1b* promoters (Higashijima et al., 1997). Tg[UAS:V2V] was generated by tagging the zebrafish *vamp2* (GenBank accession number, AAH59626) with Venus (Nagai et al., 2002) at the C-terminus, linking it to 5xUAS. Tg[egr2b-Cre] was constructed using the BAC for *egr2b* (DKEY-247F17/CR354434, Source BioScience). The Cre-polyA-Km^r^ cassette containing a Cre recombinase codon-optimized for zebrafish was introduced downstream of the *egr2b* 5’UTR. The Cre-polyA-Km^r^ cassette was amplified by PCR using pKZzCreKm plasmid as template with the primer pair egr2b-zCre-f (5’-aca caa cac att ctg tga aca tcc gag cga gtg ctt ctt agg act tca cgA TGG CTA ACT TGC TCA CTG TGC AT-3’) and Km-r (5’- aca aag cca ccg aga ctc aca ggg gct ttc tcc aaa gtt tta gct gtC ATC AAT TCA GAA GAA CTC GTC AAG AA-3’), where lower and upper cases indicate *egr2b* sequences for homologous recombination and Cre-polyA-Km^r^-annealing sequences, respectively. Tg[egr2b-Cre] was chosen from three independent BAC inserts, all of which directed recombination in r3 and r5, by crossing with Tg[zfand5b:Gal4, UAS:loxRloxG] line. For Tol2-mediated BAC transgenesis, the iTol2-amp cassette (Suster et al., 2009) was amplified by PCR with the primer pair iTol2A-f (5’-cga gcc gga agc ata aag tgt aaa gcc tgg gg tgc cta atg agt gag cta CCC TGC TCG AGC CGG GCC C-3’) and iTol2A-r (5’-ggt ttc ccg act gga aag cgg gca gtg agc gca acg caa tta atg tga gtA TTA TGA TCC TCT AGA TCA GAT CT-3’), where the lower and upper cases are pIndigoBAC-5 sequences for homologous recombination and iTol2-amp annealing sequences, respectively, was introduced into the backbone (pIndigoBAC-5, GenBank Accession; EU140754). Tg[UAS:loxRloxG] and Tg[UAS:loxGloxR] were generated by fusing the loxP-mRFP1-polyA-loxP-EGFP-polyA and loxP-EGFP-polyA-loxP-mRFP1-polyA cassettes to 5xUAS, respectively. The UAS constructs for expressing the mRFP1-tagged full-length *pcdh17*, *pcdh17-ΔCP*, *pcdh17-ΔCPT* were created by fusing these gene fragments to the *hsp70l* promoter and 4xUAS. EGFP was introduced on the other side of UAS for bidirectional induction. All transgenes are introduced into the zerbrafish genome by *Tol2*-mediated transgenesis. Gene trap lines were generated by injecting the wild type fish with the pT2GgSAIzGFFM plasmid (GA and KK, unpublished results) together with the synthesized Tol2 transposase mRNA (Abe, 2011 #72) at one-cell stage were raised and crossed with UAS:EGFP reporter line.

### Whole-mount double in situ fluorescent hybridization

Double *in situ* hybridization for *pcdh17* and EGFP were performed against Tg[mnr2b-hs:Gal4, UAS:EGFP] larvae at 40 hpf as described in (Brend and Holley, 2009). Antisense probes for *pcdh17* and EGFP were synthesized with DIG RNA labelling kit (Roche) using the coding sequence for the extracellular and transmembrane domains of *pcdh17* and the full length of EGFP as templates, respectively. The *pcdh17* fragment was cloned by PCR with the primers *pcdh17*-f1 (5’-ATG CAT CTT TCT GTT TTC CTT-3’) and *pcdh17*-r1 (5’-ACA GTT GTA TGT GCG GAT TTC-3’). Antisense probes for *pcdh17* and EGFP were synthesized with DIG RNA labeling kit (Roche) and Fluorescenin RNA labeling kit (Roche), respectively. The DIG-labeled *pcdh17* and Fluorescein-labeled EGFP were detected TSA Kit#16 AlexaFluor647 tyramide and TSA Plus Fluorescein system (NEL74100KT, PerkinElmer), respectively.

### Kaede photoconversion

Tg[mnr2b-hs:Gal4] fish was crossed with UAS:Kaede line (Scott et al., 2007) to obtain Tg[mnr2b-hs:Gal4, UAS:Kaede] embryos. mnr2b-ABNs in the Tg[mnr2b-hs:Gal4, UAS:Kaede] embryos at 44 hpf were scanned with a laser with 405 nm wave length by using Olympus FV-1000D laser confocal microscope.

### Single cell labeling

For single cell labeling of Gal4FF-expressing cells in Tg[mnr2b-hs:Gal4], ~18pg of pT2KUASGFP plasmid (Asakawa et al., 2008) was injected into a blastomere at 4- or 8-cell stage. Injected embryos were raised in the embryonic buffer containing 0.003% (w/v) *N*-Phenylthiourea (SIGMA, P7629) to inhibit melanogenesis. Only cells uniquely labeled with EGFP or clearly identifiable as single isolated cells in a hemisphere were analyzed by confocal microscopy. Soma size was defined by measuring the largest cross sectional area of the soma by ImageJ software.

### Inverse PCR and RT-PCR

The flanking genomic sequences for Tg[gSAIzGFFM773B] and Tg[gSAIzGFFM1068A] were cloned by the inverse PCR performed as described previously (Urasaki et al., 2006). For RT-PCR, the total RNA was prepared from each of fifty Tg[gSAIzGFFM773B] and Tg[gSAIzGFFM1068A] larvae by homogenizing them in 1 ml of Trizol Reagent (Life Technologies). RT-PCR was performed by using the following six primers as indicated in the SupFig1, tnnt3b-f (5’-gat gga gat aag gtg gac ttt gac-3’), tnnt3b-r (5’-ttt act gag ctc ctc cac tct ctt-3’), tnnt1-f (5’-ggt gag agg gta gat ttt gat gat-3’), tnnt1-r (5’-cca aga ggt caa att tct cag act-3’), Gal4FF-f (5’-tgc tga gct cta tcg agc agg-3’) and Gal4FF-r (5’-ggc agc ata tcc aga tcg aag-3’). The insert Tg[gSAIzGFFM773B] was located in the sixth intron of *tnnt1* (GenBank: AF425255.1). The insert Tg[gSAIzGFFM1068A] was integrated in an internal intron of *tnnt3b* (GenBank: AF512526.1).

### Generation of zebrafish mutants by CRISPR-Cas9 method

For the generation of *sall4* mutant fish, eight target sites for CRISPR-Cas9-mediated cleavage were selected in the middle region of the 2^nd^ exon based on CRISPRscan (Moreno-Mateos et al., 2015). The guide RNAs (gRNAs) were produced by *in vitro* transcription by MEGAshortscript™ T7 Transcription Kit (Thermo Fisher Scientific, AM1354) by using PCR-amplified DNA templates. hSpCas9 was *in vitro* transcribed with mMESSAGE mMACHINE Kit (Thermo Fisher Scientific, AM1340) by using pCS2+hSpCas9 plasmid as a template (a gift from Masato Kinoshita, Addgene plasmid # 51815). Wild type embryos were injected at one-cell stage with 25 pg of sgRNA and 300 pg of hSpCas9 mRNA. The sgRNA that yielded the highest cleavage activity in the heteroduplex mobility assay (HMA) (target sequence: GGGGAAGCAGCGCTGGAAGTcgg, where the protospacer adjacent motif (PAM) was indicated by lower cases) was used for the generation of *sall4-d11* allele. For the generation of *pcdh17* mutant fish, nine target sites were selected in either the extracellular or cytoplasmic domain-coding exon. The *pcdh17-d77* allele was generated with the Alt-R® CRISPR-Cas9 system (IDT) using a crRNA containing TAGCGGTGCGTCAGGGGG as a target sequence followed by the PAM, CGG.

### Statistics

The difference in fluorescence intensities (Fig.1 J) was evaluated by unpaired t test (two-tailed). For td-Tomato signal, p = 0.12, t = 1.63, df = 18 and N = 10 neuromuscular junctions from 5 animals. For EGFP signal, p = 0.02, t = 2.49, df = 16 and N = 10. The difference in soma size (Fig.2 G) was evaluated by unpaired t test (two-tailed). For r5, p < 0.0001, t = 4.99, df = 18.91 and N = 12 cells for each of dorsal and ventral clusters. For r6, N = 12, p = 0.009, t = 4.19, df = 13.74 and N = 10 and 7 cells for dorsal and ventral clusters, respectively. Difference of the dorsal indices between wild type and the mutant condition was evaluated by unpaired t test (two-tailed). For pcdh17-d77 heterozygotes, p = 0.149, t = 1.49, df = 25.56 and N = 10 and 18 hemibrains from 5 and 9 animals, respectively. For *pcdh17-d77* homozygotes, p = 0.014, t = 2.71, df = 17.66 and N = 10 hemibrains from 5 animals for each of wild type and *pcdh17-d77* homozygotes. Statistical analyses were performed by using Prism 6 software.

### Microscopic analysis

A fluorescence stereomicroscope (MZ16FA, Leica) equipped with a CCD camera (DFC300FX, Leica) was used to observe and take images of GFP-expressing embryos. For confocal microscopy, a live embryo or larva was embedded in 1 % low-melting agarose (NuSieve® GTG® Agarose, Lonza) on a Glass Base dish (IWAKI, 3010-035) and subject to confocal microscopy using Olympus FV-1000D laser confocal microscope. Images of live embryos and larvae were acquired as serial sections along the z-axis and processed with Olympus Fluoview Ver2.1b Viewer and Adobe Photoshop CS6. Fluorescent intensity measurements were performed using imageJ software.

### Data availability

The data that support the findings in this study are available within the article and its Supplementary Information files, and from the corresponding authors upon request.

## References

Al-Baradie, R., Yamada, K., St Hilaire, C., Chan, W.M., Andrews, C., McIntosh, N., Nakano, M., Martonyi, E.J., Raymond, W.R., Okumura, S., et al. (2002). Duane radial ray syndrome (Okihiro syndrome) maps to 20q13 and results from mutations in SALL4, a new member of the SAL family. Am J Hum Genet 71, 1195–1199.

Ampatzis, K., Song, J., Ausborn, J., and El Manira, A. (2014). Separate microcircuit modules of distinct v2a interneurons and motoneurons control the speed of locomotion. Neuron 83, 934–943.

Asakawa, K., Abe, G., and Kawakami, K. (2013). Cellular dissection of the spinal cord motor column by BAC transgenesis and gene trapping in zebrafish. Front Neural Circuits 7, 100.

Asakawa, K., Higashijima, S., and Kawakami, K. (2012). An mnr2b/hlxb9lb enhancer trap line that labels spinal and abducens motor neurons in zebrafish. Dev Dyn 241, 327–332.

Asakawa, K., and Kawakami, K. (2009). The Tol2-mediated Gal4-UAS method for gene and enhancer trapping in zebrafish. Methods 49, 275–281.

Asakawa, K., Suster, M.L., Mizusawa, K., Nagayoshi, S., Kotani, T., Urasaki, A., Kishimoto, Y., Hibi, M., and Kawakami, K. (2008). Genetic dissection of neural circuits by Tol2 transposon-mediated Gal4 gene and enhancer trapping in zebrafish. Proc Natl Acad Sci U S A 105, 1255–1260.

Astick, M., Tubby, K., Mubarak, W.M., Guthrie, S., and Price, S.R. (2014). Central topography of cranial motor nuclei controlled by differential cadherin expression. Curr Biol 24, 2541–2547.

Biswas, S., and Jontes, J.D. (2009). Cloning and characterization of zebrafish protocadherin-17. Dev Genes Evol 219, 265–271.

Brend, T., and Holley, S.A. (2009). Zebrafish whole mount high-resolution double fluorescent in situ hybridization. J Vis Exp.

Buttner-Ennever, J.A., Horn, A.K., Scherberger, H., and D’Ascanio, P. (2001). Motoneurons of twitch and nontwitch extraocular muscle fibers in the abducens, trochlear, and oculomotor nuclei of monkeys. J Comp Neurol 438, 318–335.

Cabrera, B., Pasaro, R., and Delgado-Garcia, J.M. (1989). Cytoarchitectonic organisation of the abducens nucleus in the pigeon (Columbia livia). J Anat 166, 203–211.

Cabrera, B., Torres, B., Pasaro, R., Pastor, A.M., and Delgado-Garcia, J.M. (1992). A morphological study of abducens nucleus motoneurons and internuclear neurons in the goldfish (Carassius auratus). Brain Res Bull 28, 137–144.

Chilton, J.K., and Guthrie, S. (2016). Axons get ahead: Insights into axon guidance and Congenital Cranial Dysinnervation Disorders. Dev Neurobiol.

Clark, C., Austen, O., Poparic, I., and Guthrie, S. (2013). alpha2-Chimaerin regulates a key axon guidance transition during development of the oculomotor projection. J Neurosci 33, 16540–16551.

da Silva Costa, R.M., Kung, J., Poukens, V., Yoo, L., Tychsen, L., and Demer, J.L. (2011). Intramuscular innervation of primate extraocular muscles: unique compartmentalization in horizontal recti. Invest Ophthalmol Vis Sci 52, 2830–2836.

Demer, J.L., Clark, R.A., Lim, K.H., and Engle, E.C. (2007). Magnetic resonance imaging of innervational and extraocular muscle abnormalities in Duane-radial ray syndrome. Invest Ophthalmol Vis Sci 48, 5505–5511.

Eberhorn, A.C., Ardeleanu, P., Buttner-Ennever, J.A., and Horn, A.K. (2005). Histochemical differences between motoneurons supplying multiply and singly innervated extraocular muscle fibers. J Comp Neurol 491, 352–366.

Eberhorn, A.C., Buttner-Ennever, J.A., and Horn, A.K. (2006). Identification of motoneurons supplying multiply- or singly-innervated extraocular muscle fibers in the rat. Neuroscience 137, 891–903.

Epley, K.E., Urban, J.M., Ikenaga, T., and Ono, F. (2008). A modified acetylcholine receptor delta-subunit enables a null mutant to survive beyond sexual maturation. J Neurosci 28, 13223–13231.

Ferrario, J.E., Baskaran, P., Clark, C., Hendry, A., Lerner, O., Hintze, M., Allen, J., Chilton, J.K., and Guthrie, S. (2012). Axon guidance in the developing ocular motor system and Duane retraction syndrome depends on Semaphorin signaling via alpha2-chimaerin. Proc Natl Acad Sci U S A 109, 14669–14674.

Fuchs, A.F., and Luschei, E.S. (1970). Firing patterns of abducens neurons of alert monkeys in relationship to horizontal eye movement. J Neurophysiol 33, 382–392.

Gaufo, G.O., Thomas, K.R., and Capecchi, M.R. (2003). Hox3 genes coordinate mechanisms of genetic suppression and activation in the generation of branchial and somatic motoneurons. Development 130, 5191–5201.

Gestrin, P., and Sterling, P. (1977). Anatomy and physiology of goldfish oculomotor system. II. Firing patterns of neurons in abducens nucleus and surrounding medulla and their relation to eye movements. J Neurophysiol 40, 573–588.

Gilland, E., and Baker, R. (2005). Evolutionary patterns of cranial nerve efferent nuclei in vertebrates. Brain Behav Evol 66, 234–254.

Greaney, M.R., Privorotskiy, A.E., D’Elia, K.P., and Schoppik, D. (2017). Extraocular motoneuron pools develop along a dorsoventral axis in zebrafish, Danio rerio. J Comp Neurol 525, 65–78.

Guthrie, S. (2007). Patterning and axon guidance of cranial motor neurons. Nat Rev Neurosci 8, 859–871.

Hayashi, S., Inoue, Y., Kiyonari, H., Abe, T., Misaki, K., Moriguchi, H., Tanaka, Y., and Takeichi, M. (2014). Protocadherin-17 mediates collective axon extension by recruiting actin regulator complexes to interaxonal contacts. Dev Cell 30, 673–687.

Higashijima, S., Hotta, Y., and Okamoto, H. (2000). Visualization of cranial motor neurons in live transgenic zebrafish expressing green fluorescent protein under the control of the islet-1 promoter/enhancer. J Neurosci 20, 206–218.

Higashijima, S., Okamoto, H., Ueno, N., Hotta, Y., and Eguchi, G. (1997). High-frequency generation of transgenic zebrafish which reliably express GFP in whole muscles or the whole body by using promoters of zebrafish origin. Dev Biol 192, 289–299.

Holmgren, N. (1922). Points of view concerning forebrain morphology in lower vertebrates. J Comp Neurol 34, 34.

Hoshina, N., Tanimura, A., Yamasaki, M., Inoue, T., Fukabori, R., Kuroda, T., Yokoyama, K., Tezuka, T., Sagara, H., Hirano, S., et al. (2013). Protocadherin 17 regulates presynaptic assembly in topographic corticobasal Ganglia circuits. Neuron 78, 839–854.

Kaminski, H.J., Richmonds, C.R., Kusner, L.L., and Mitsumoto, H. (2002). Differential susceptibility of the ocular motor system to disease. Ann N Y Acad Sci 956, 42–54.

Kasprick, D.S., Kish, P.E., Junttila, T.L., Ward, L.A., Bohnsack, B.L., and Kahana, A. (2011). Microanatomy of adult zebrafish extraocular muscles. PLoS One 6, e27095.

Khanna, S., Cheng, G., Gong, B., Mustari, M.J., and Porter, J.D. (2004). Genome-wide transcriptional profiles are consistent with functional specialization of the extraocular muscle layers. Invest Ophthalmol Vis Sci 45, 3055–3066.

Kohlhase, J., Heinrich, M., Schubert, L., Liebers, M., Kispert, A., Laccone, F., Turnpenny, P., Winter, R.M., and Reardon, W. (2002). Okihiro syndrome is caused by SALL4 mutations. Hum Mol Genet 11, 2979–2987.

Lefebvre, J.L., Kostadinov, D., Chen, W.V., Maniatis, T., and Sanes, J.R. (2012). Protocadherins mediate dendritic self-avoidance in the mammalian nervous system. Nature 488, 517–521.

Liu, Q., Bhattarai, S., Wang, N., and Sochacka-Marlowe, A. (2015). Differential expression of protocadherin-19, protocadherin-17, and cadherin-6 in adult zebrafish brain. J Comp Neurol 523, 1419–1442.

Liu, Q., Chen, Y., Pan, J.J., and Murakami, T. (2009). Expression of protocadherin-9 and protocadherin-17 in the nervous system of the embryonic zebrafish. Gene Expr Patterns 9, 490–496.

Lu, C.C., Appler, J.M., Houseman, E.A., and Goodrich, L.V. (2011). Developmental profiling of spiral ganglion neurons reveals insights into auditory circuit assembly. J Neurosci 31, 10903–10918.

Lu, Q.R., Sun, T., Zhu, Z., Ma, N., Garcia, M., Stiles, C.D., and Rowitch, D.H. (2002). Common developmental requirement for Olig function indicates a motor neuron/oligodendrocyte connection. Cell 109, 75–86.

Lumsden, A., and Keynes, R. (1989). Segmental patterns of neuronal development in the chick hindbrain. Nature 337, 424–428.

Lyons, P.J., Ma, L.H., Baker, R., and Fricker, L.D. (2010). Carboxypeptidase A6 in zebrafish development and implications for VIth cranial nerve pathfinding. PLoS One 5, e12967.

Ma, L.H., Grove, C.L., and Baker, R. (2014). Development of oculomotor circuitry independent of hox3 genes. Nat Commun 5, 4221.

Manzanares, M., Cordes, S., Kwan, C.T., Sham, M.H., Barsh, G.S., and Krumlauf, R. (1997). Segmental regulation of Hoxb-3 by kreisler. Nature 387, 191–195.

McKay, I.J., Lewis, J., and Lumsden, A. (1997). Organization and development of facial motor neurons in the kreisler mutant mouse. Eur J Neurosci 9, 1499–1506.

McKay, I.J., Muchamore, I., Krumlauf, R., Maden, M., Lumsden, A., and Lewis, J. (1994). The kreisler mouse: a hindbrain segmentation mutant that lacks two rhombomeres. Development 120, 2199–2211.

McLean, D.L., Fan, J., Higashijima, S., Hale, M.E., and Fetcho, J.R. (2007). A topographic map of recruitment in spinal cord. Nature 446, 71–75.

Menelaou, E., VanDunk, C., and McLean, D.L. (2014). Differences in the morphology of spinal V2a neurons reflect their recruitment order during swimming in larval zebrafish. J Comp Neurol 522, 1232–1248.

Miyake, N., Chilton, J., Psatha, M., Cheng, L., Andrews, C., Chan, W.M., Law, K., Crosier, M., Lindsay, S., Cheung, M., et al. (2008). Human CHN1 mutations hyperactivate alpha2-chimaerin and cause Duane’s retraction syndrome. Science 321, 839–843.

Moens, C.B., Yan, Y.L., Appel, B., Force, A.G., and Kimmel, C.B. (1996). valentino: a zebrafish gene required for normal hindbrain segmentation. Development 122, 3981–3990.

Moreno-Mateos, M.A., Vejnar, C.E., Beaudoin, J.D., Fernandez, J.P., Mis, E.K., Khokha, M.K., and Giraldez, A.J. (2015). CRISPRscan: designing highly efficient sgRNAs for CRISPR-Cas9 targeting in vivo. Nat Methods 12, 982–988.

Morgan, D.L., and Proske, U. (1984). Vertebrate slow muscle: its structure, pattern of innervation, and mechanical properties. Physiol Rev 64, 103–169.

Mountoufaris, G., Chen, W.V., Hirabayashi, Y., O’Keeffe, S., Chevee, M., Nwakeze, C.L., Polleux, F., and Maniatis, T. (2017). Multicluster Pcdh diversity is required for mouse olfactory neural circuit assembly. Science 356, 411–414.

Nagai, T., Ibata, K., Park, E.S., Kubota, M., Mikoshiba, K., and Miyawaki, A. (2002). A variant of yellow fluorescent protein with fast and efficient maturation for cell-biological applications. Nat Biotechnol 20, 87–90.

Nugent, A.A., Park, J.G., Wei, Y., Tenney, A.P., Gilette, N.M., DeLisle, M.M., Chan, W.M., Cheng, L., and Engle, E.C. (2017). Mutant alpha2-chimaerin signals via bidirectional ephrin pathways in Duane retraction syndrome. J Clin Invest 127, 1664–1682.

Orr, H. (1887). Contribution to the embryology of the lizard; With especial reference to the central nervous system and some organs of the head; together with observations on the origin of the vertebrates. Journal of Morphology 1, 311–372.

Park, J.G., Tischfield, M.A., Nugent, A.A., Cheng, L., Di Gioia, S.A., Chan, W.M., Maconachie, G., Bosley, T.M., Summers, C.G., Hunter, D.G., et al. (2016). Loss of MAFB Function in Humans and Mice Causes Duane Syndrome, Aberrant Extraocular Muscle Innervation, and Inner-Ear Defects. Am J Hum Genet 98, 1220–1227.

Pastor, A.M., Torres, B., Delgado-Garcia, J.M., and Baker, R. (1991). Discharge characteristics of medial rectus and abducens motoneurons in the goldfish. J Neurophysiol 66, 2125–2140.

Peng, M., Poukens, V., da Silva Costa, R.M., Yoo, L., Tychsen, L., and Demer, J.L. (2010). Compartmentalized innervation of primate lateral rectus muscle. Invest Ophthalmol Vis Sci 51, 4612–4617.

Prince, V.E., Moens, C.B., Kimmel, C.B., and Ho, R.K. (1998). Zebrafish hox genes: expression in the hindbrain region of wild-type and mutants of the segmentation gene, valentino. Development 125, 393–406.

Robinson, D.A. (1970). Oculomotor unit behavior in the monkey. J Neurophysiol 33, 393–403.

Sakaki-Yumoto, M., Kobayashi, C., Sato, A., Fujimura, S., Matsumoto, Y., Takasato, M., Kodama, T., Aburatani, H., Asashima, M., Yoshida, N., et al. (2006). The murine homolog of SALL4, a causative gene in Okihiro syndrome, is essential for embryonic stem cell proliferation, and cooperates with Sall1 in anorectal, heart, brain and kidney development. Development 133, 3005–3013.

Scott, E.K., Mason, L., Arrenberg, A.B., Ziv, L., Gosse, N.J., Xiao, T., Chi, N.C., Asakawa, K., Kawakami, K., and Baier, H. (2007). Targeting neural circuitry in zebrafish using GAL4 enhancer trapping. Nat Methods 4, 323–326.

Siebeck, R., and Kruger, P. (1955). [The histological structure of the extrinsic ocular muscles as an indication of their function]. Albrecht Von Graefes Arch Ophthalmol 156, 636–652.

Spencer, R.F., and Porter, J.D. (1988). Structural organization of the extraocular muscles. Rev Oculomot Res 2, 33–79.

Spencer, R.F., and Porter, J.D. (2006). Biological organization of the extraocular muscles. Prog Brain Res 151, 43–80.

Sterling, P., and Gestrin, P. (1975). Goldfish abducens motoneurons: physiological and anatomical specialization. Science 189, 1091–1093.

Suster, M.L., Sumiyama, K., and Kawakami, K. (2009). Transposon-mediated BAC transgenesis in zebrafish and mice. BMC Genomics 10, 477.

Thaler, J., Harrison, K., Sharma, K., Lettieri, K., Kehrl, J., and Pfaff, S.L. (1999). Active suppression of interneuron programs within developing motor neurons revealed by analysis of homeodomain factor HB9. Neuron 23, 675–687.

Tischfield, M.A., Bosley, T.M., Salih, M.A., Alorainy, I.A., Sener, E.C., Nester, M.J., Oystreck, D.T., Chan, W.M., Andrews, C., Erickson, R.P., et al. (2005). Homozygous HOXA1 mutations disrupt human brainstem, inner ear, cardiovascular and cognitive development. Nat Genet 37, 1035–1037.

Urasaki, A., Morvan, G., and Kawakami, K. (2006). Functional dissection of the Tol2 transposable element identified the minimal cis-sequence and a highly repetitive sequence in the subterminal region essential for transposition. Genetics 174, 639–649.

Vaage, S. (1969). The segmentation of the primitive neural tube in chick embryos (Gallus domesticus). A morphological, histochemical and autoradiographical investigation. Ergeb Anat Entwicklungsgesch 41, 3–87.

Whitman, M.C., and Engle, E.C. (2017). Ocular Congenital Cranial Dysinnervation Disorders (CCDDs): Insights into Axon Growth and Guidance. Hum Mol Genet.

Yu Wai Man, C.Y., Chinnery, P.F., and Griffiths, P.G. (2005). Extraocular muscles have fundamentally distinct properties that make them selectively vulnerable to certain disorders. Neuromuscul Disord 15, 17–23.

Zannino, D.A., and Appel, B. (2009). Olig2+ precursors produce abducens motor neurons and oligodendrocytes in the zebrafish hindbrain. J Neurosci 29, 2322–2333.

Zhou, Q., and Anderson, D.J. (2002). The bHLH transcription factors OLIG2 and OLIG1 couple neuronal and glial subtype specification. Cell 109, 61–73.

