## Supplementary Materials for "Protocadherin-mediated cell repulsion controls the central topography and efferent projections of the abducens nucleus"

**Supplementary information**

**Supplementary Fig 1. Compartmental organization of the lateral rectus muscle characterized by troponins.**

(A) Dorsal view of the left lateral rectus muscle in a Tg[gSAIzGFFM1068A, UAS:EGFP, actc1b:tdT-chrnd] larva at 5dpf. The double arrowheads indicate the presumptive initial contact sites between the abducens motor neurons and lateral rectus muscle. Rostral is to the top. (B) Tg[gSAIzGFFM1068A], indicated as M1068A, traps *tnnt3b*. (C) Dorsal view of the left lateral rectus muscle in a Tg[gSAIzGFF773B, UAS:EGFP, actc1b:tdT-chrnd] larva at 5dpf. (D) Tg[gSAIzGFF773B], indicated as M773B, traps *tnnt1*. The open boxes represent exons of the troponin genes. Chimeric transcript between the troponin genes and Gal4 was detectable by RT-PCR (bottom). The filled and open arrowheads show the position and direction (f: forward, and r: reverse) of primers annealing with the troponin exons and Gal4, respectively. For each set of primer pairs, cDNA reverse-transcribed from poly A-tailed RNA of fish with (G, left lane) or without (-, right lane) the gene trap construct was examined. The ubiquitously expressed *zfand5b* was used as a control. The bars indicate 50 $\mu$m in A, C and 100 bp in B, D.

**Supplementary Fig 2. A descending interneuron labeled by Tg[mnr2b-hs:Gal4].**

Sparse cell labeling identified a descending interneuron in the dorsal side of the r5 in Tg[mnr2b-hs:Gal4] larvae at 5 dpf. The vertical dashed line shows the position of the midline. The bar indicates 50 $\mu$m.

**Supplementary Fig 3. Tg[mnr2b-GFP] line marks mnr2b-ABNs.**

The dorsal (A-C) and ventral (D-F) sides of r5 and r6 in Tg[mnr2b-GFP, mnr2b-hs:Gal4, UAS:mCherry] larva at 80 hpf. The vertical dashed lines show the position of the midline. (G-I) Tg[mnr2b-GFP] also labels the spinal motor neurons expressing Gal4 form Tg[mnr2b-hs:Gal4], which are labeled with UAS:mCherry (magenta). The horizontal dashed lines demarcate the dorsal and ventral limits of the spinal cord. The bar indicates 50 $\mu$m.

**Supplementary Fig 4.** *sall4* knockout fish forms the neuromuscular connection between mnr2b-ABN and lateral rectus muscle.

(A) The structure of zebrafish and human Sall4/SALL4 protein. Zebrafish Sall4 consists of 1091 amino acid. The *sall4-d11* allele encodes the N-terminus third of Sall4 followed by 15 amino-acid peptide. Human SALL4 (bottom) has similar molecular structure with the zebrafish orthologue. The asterisks indicate the approximate positions of deletion and insertion mutations causing protein truncation in Okihiro/DRRS syndrome patiens. The orange ovals represent the zinc finger domains. (B) The 11-bp deletion in the *sall4-d11* mutant allele (red) generates a premature stop codon. The nucleotides at the position from +922 to +932, where A in the initiation codon is +1, are deleted in *sall4-d11*. The premature stop codon is shown in green. The underlined CCG (CGG in the opposite strand) represents the position of the protospacer adjacent motif (PAM) for the sgRNA for Cas9-mediated cleavage. (C, D) The dorsal (top) and lateral (bottom) views of the wild type (C) and *sall4-d11* (D) larvae at 5 dpf. The larva homozygous for *sall4-d11* shows morphological abnormalities. The arrow indicates the cardiac edema (D, bottom). Arrowheads show the eye edema (inset). (E-J) The dorsal view of the lateral rectus muscle (E, H) and the ventral (F, I) and dorsal cluster (G, J) of abducens nuclei in the wild type (left panels) and the *sall4-d11* homozygote (right panels) with Tg[mnr2b-hs:Gal4, UAS:GFP, actc1b-hs:tdT-chrnd] background at 5dpf. The double arrowheads indicate CEPZ. The white and black arrowheads indicate the position of r5 and r6, respectively. The brackets indicate the dorsal cluster. The dashed lines show the midline. The bars indicate 50 $\mu$m.

**Supplementary Fig 5. Embryonic and larval phenotypes occurring from *pcdh17-d77* mutant crosses.**

(A) Incidence of developmental defects occurring during the embryonic (red) and larval (larval) stages observed among the offspring from indicated parental fish. The wild type and *pcdh17-d77* alleles were indicated as + and *d77*, respectively. Embryonic phenotypes typically include shortened body due to defective gastrulation, which occasionally resulted in embryonic death due to yolk damage. Either of failure in forming a fully inflated swim bladder on 5 dpf or morphological abnormalities such as cardiac edema was considered as larval defects. Fish that initiated free swimming with a fully inflated swim bladder on 5 dpf were categorized as normal (green). In total, 528, 322, 185 fish were examined from *pcdh17-d77*/+ incross (left, 9 crosses from 5 independent parental pairs), female *pcdh17-d77*/+ outcross (middle, 8 crosses from 5 independent parental pairs), and male *pcdh17-d77*/+ outcross (right, 4 crosses from 4 independent parental pairs), respectively. (B) Examples of the embryonic defect observed at 20 hpf.

**Supplementary Fig 6. Misguidance and defasciculation phenotypes produced by overexpressing Pcdh17-FL-mRFP1.**

(A) Dorsal view of the normal mnr2b-ABN projection in a Tg[mnr2b-hs:Gal4, EGFP:UAS:pcdh17-FL-mRFP1] larva at 3 dpf. (B) Misguidance of the mnr2b-ABNs in Tg[mnr2b-hs:Gal4, EGFP:UAS:pcdh17-FL-mRFP1] larva. Yellow arrows indicate the misguided axons. (C) Defasciculation of the mnr2b-ABN axon in Tg[mnr2b-hs:Gal4, EGFP:UAS:pcdh17-FL-mRFP1] larva. White arrows indicate the defasciculated axons. The double arrowheads indicate CEPZ. The white and black arrowheads indicate the position of r5 and r6, respectively. The bars indicate 50 $\mu$m in A, B and 20 $\mu$m in C.

**Supplementary Fig 7. Deletion of the transmenbrane domain from *pcdh17-ΔCP* abolished the clumping effect.**

(B) Schematic drawing of UAS constructs for pcdh17-ΔCP and pcdh17-ΔCPT expression. EC and TM stand for extracellular and transmembrane domains, respectively. EGFP and the C-terminally mRFP1-tagged *pcdh17* genes were place on each side of 4xUAS. The *hsp70l* promoter was used as a basal promoter for the *pcdh17* genes. (B-F) Dorsal and rear views of r5 and r6 in 3 dpf larvae carrying

Tg[mnr2b-hs:Gal4] and the transgene indicated as above. The white and black arrowheads indicate the position of r5 and r6, respectively. (G) Dorsal view of the caudal brain. The double arrowheads indicate the presumptive initial contact sites with the lateral rectus muscle. (H) Frequency of abnormal axon growth and fasciculation. The numbers of the nerves investigated are shown in the bar. NA, not applicable due to severe axon growth defect. The panels B and C show the same fish as in Fig. 5F. In H, the data for EGFP and pcdh17-ΔCP in Fig 5. I and J were used for reference. The horizontal scale bars indicate 20 $\mu$m in B and 50 $\mu$m in G.

**Supplementary Fig 8. Clumping of axons and somas of the spinal motor neurons expressing *pcdh17-ΔCP-mRFP1.***

The lateral views of the trunk region of Tg[mnr2b-hs:Gal4, UAS:EGFP] (A, B) and Tg[mnr2b-hs:Gal4, EGFP:UAS: pcdh17-ΔCP-mRFP1] (C, D) larvae at 3 dpf. The dashed lines indicate the ventral limit of the spinal cord. The bar indicates 100 $\mu$m.

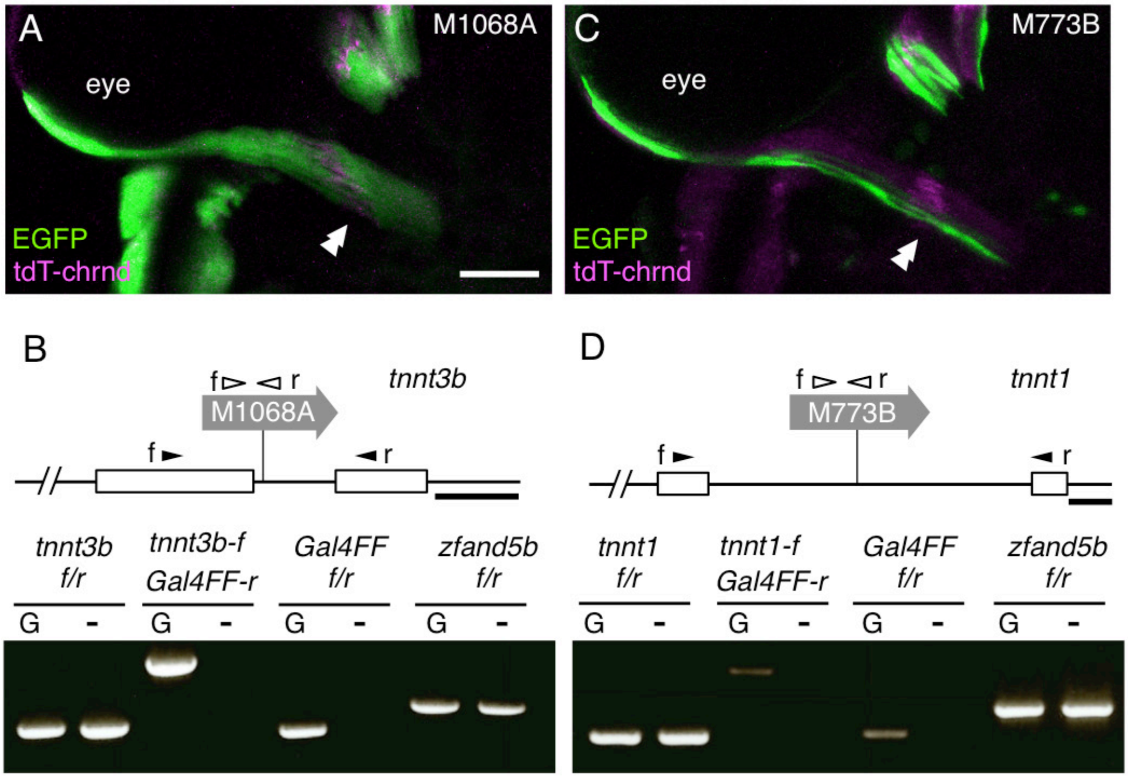


Sup Fig 1, Asakawa et al.

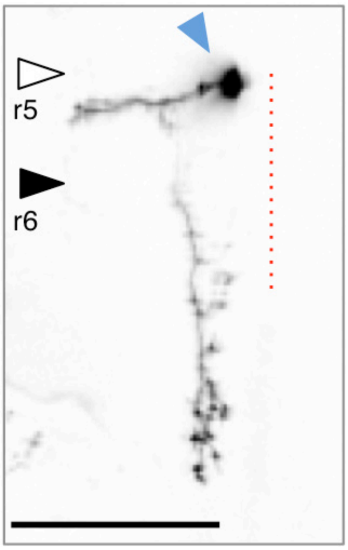


Sup. Figure 2,  Asakawa et al.

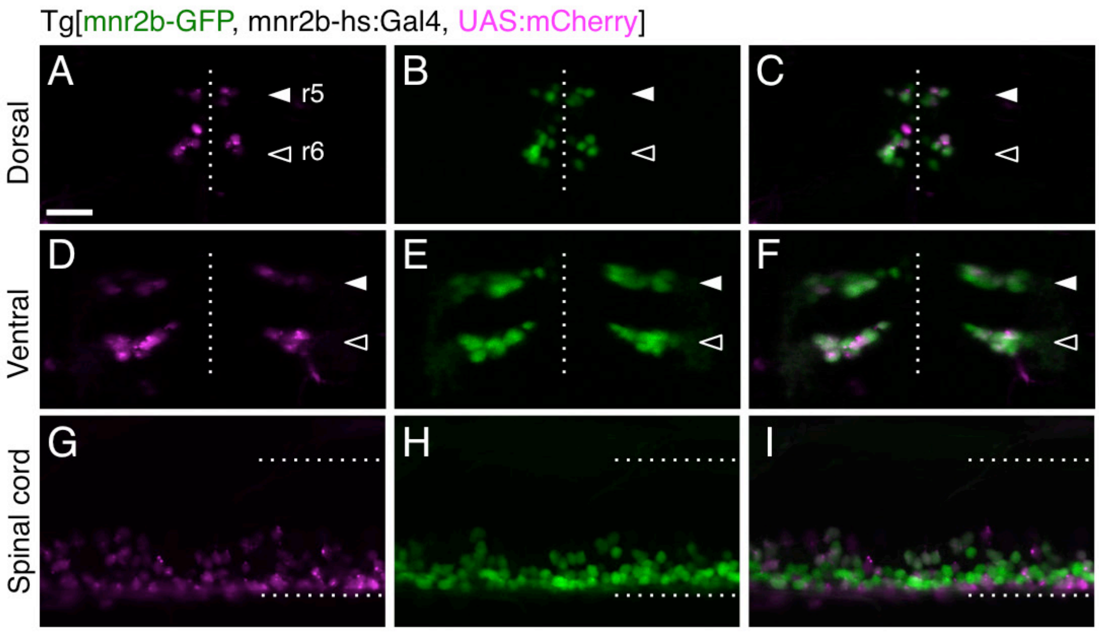


Sup. Figure 3,  Asakawa et al.

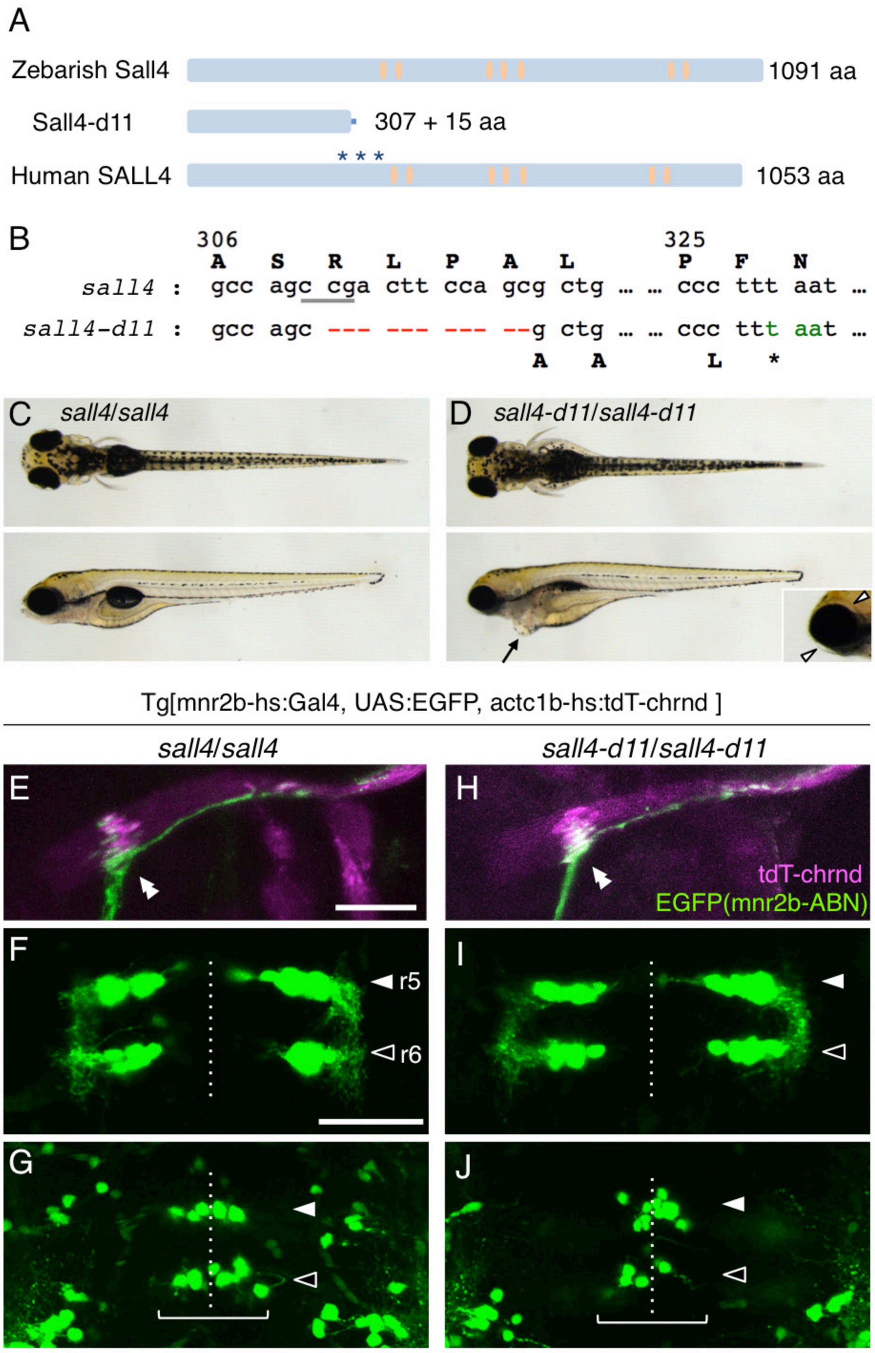


Sup Fig 4, Asakawa et al.

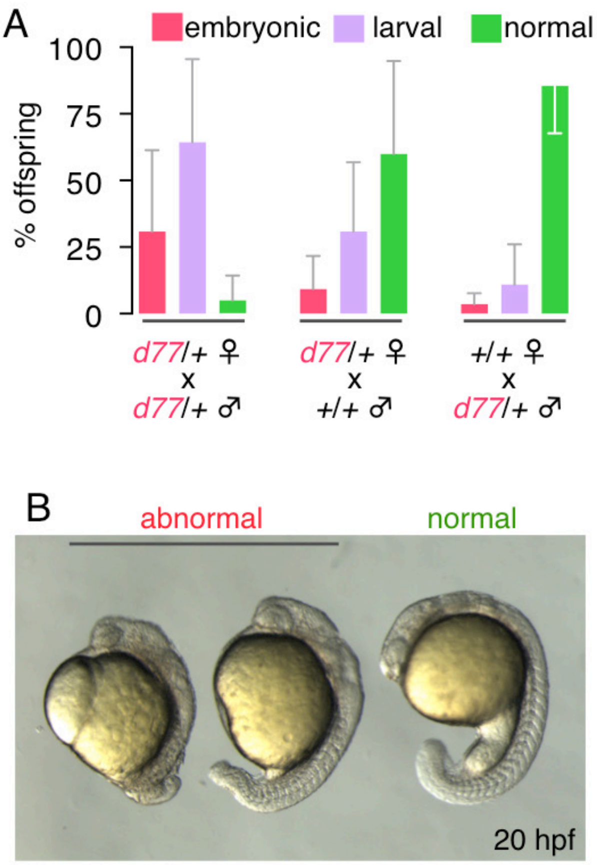


Sup. Figure 5, Asakawa et al.

Tg[mnr2b-hs:Gal4, EGFP:UAS:pcdh17-FL-mRFP1]

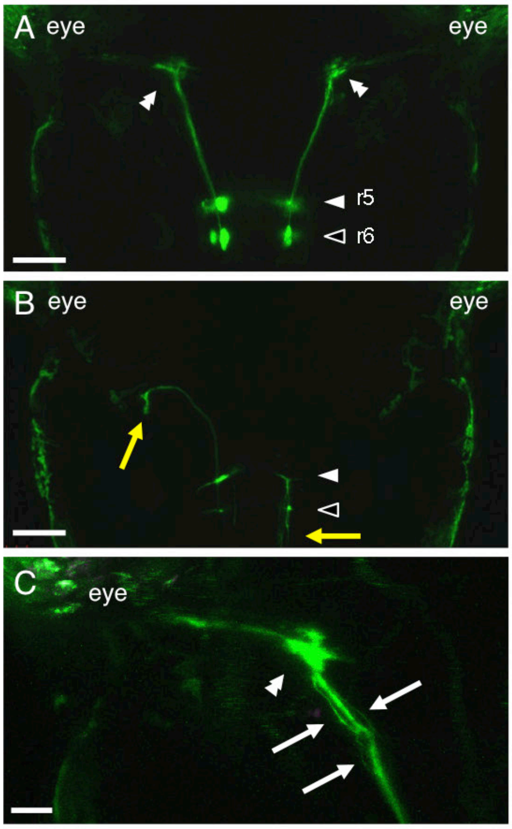


Sup. Figure 6, Asakawa et al.

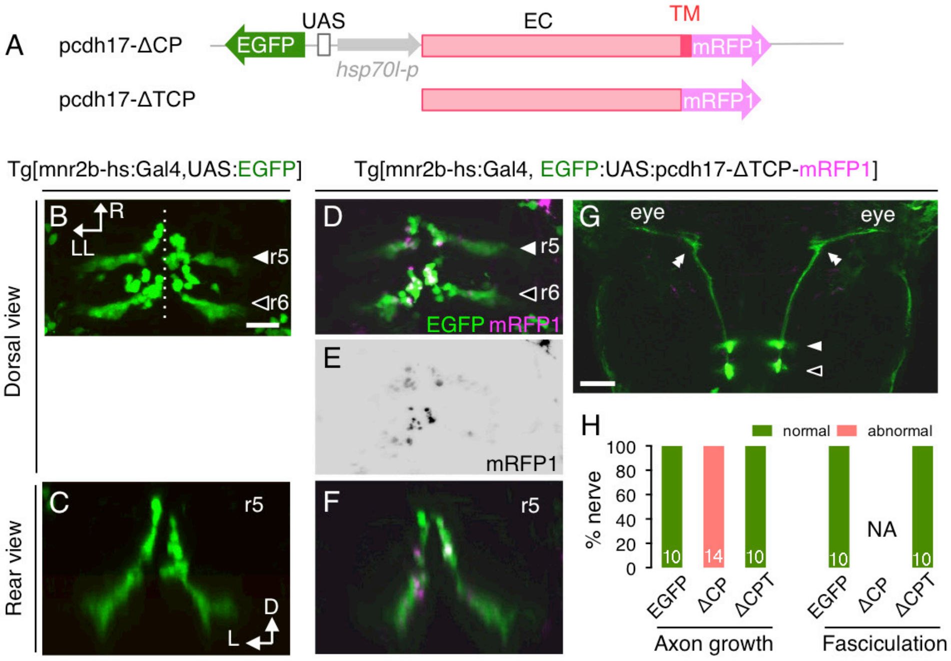


Sup. Figure 7, Asakawa et al.

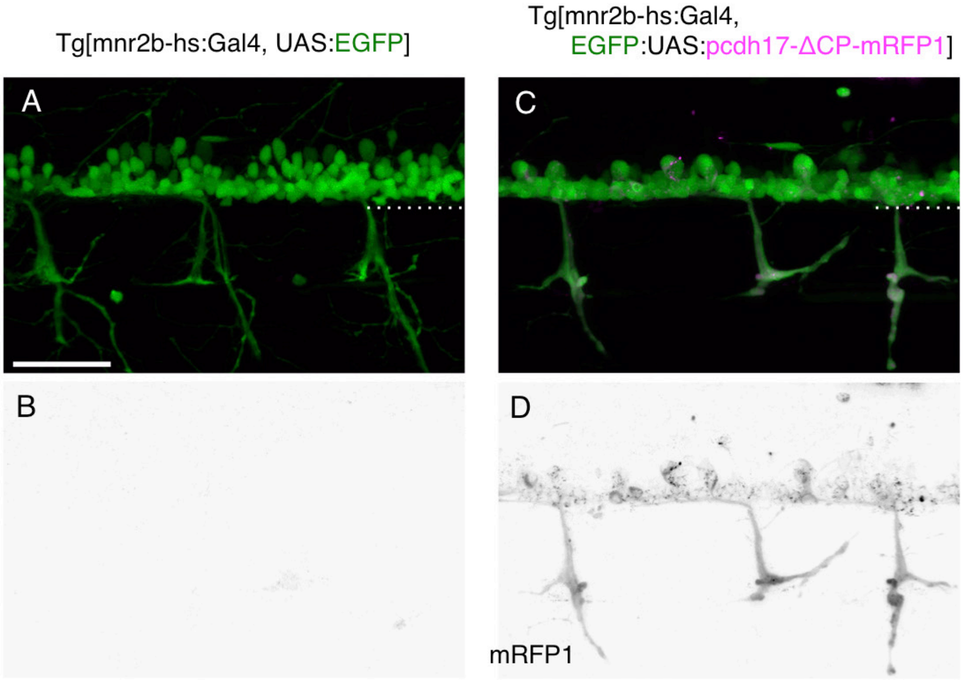


Sup. Figure 8, Asakawa et al.
